# Aridification and habitat shifts are associated with diversification in Australian diplodactylid geckos

**DOI:** 10.64898/2026.03.23.713808

**Authors:** Sarin Tiatragul, Ian G. Brennan, Alexander Skeels, Damien Esquerré, Stephen M. Zozaya, J. Scott Keogh, Mitzy Pepper

**Affiliations:** Division of Ecology & Evolution, Research School of Biology, The Australian National University, Canberra, ACT 2601, Australia; Biodiversity and Geosciences, Queensland Museum, PO Box 3300, South Brisbane BC, Queensland 4101, Australia; Environmental Futures Research Centre, School of Science, University of Wollongong, Wollongong, NSW 2500, Australia

**Keywords:** diversification, functional diversity, squamates, trait evolution, biogeography, Eocene Oligocene Transition (EOT), Oligocene Drowning, Zealandia

## Abstract

Continental radiations record the long-term interplay between environmental change, ecological opportunity, and lineage diversification across large geographic scales. The gecko family Diplodactylidae represents one such radiation with ∼200 species distributed across Australia, New Caledonia, and Aotearoa New Zealand, occupying ecological forms ranging from burrow-dwelling desert specialists to canopy climbers, and diversifying over a ∼45 Ma history shaped by dramatic continental environmental change. Using ∼5000 nuclear loci, we reconstructed phylogenetic relationships and divergence times, estimated ancestral ecology and biomes using newly generated biome-through-time maps for Australia, and modeled the effects of ecological states on diversification and morphology. Crown diplodactylids originated in the mid-Eocene (∼45 Ma), with the core Australian clade radiating in the Oligocene (∼28 Ma), substantially younger than previous estimates. Ancestral state estimation indicated arboreal origins in mesic environments, followed by repeated transitions into open habitats and expansion into semi-arid and arid biomes. Despite moderate differences in diversification rates among ecological states, rate variation was better explained by unmeasured hidden states than by observed ecological states, suggesting that our broad ecological categorization alone does not explain diversification dynamics within this radiation. Size varies with ecological state, while tail length shows no detectable ecological association and remains phylogenetically conserved. These patterns indicate that environmental change and biome transformation were associated with ecological opportunity, coinciding with diversification through repeated habitat transitions and morphological divergence, providing a macroevolutionary framework linking environmental change, ecological expansion, and trait evolution in a continental radiation.

---

Continental radiations provide powerful systems for examining how ecological opportunity, environmental change, and morphological diversification interact over deep time. Key questions in such systems concern how ecological transitions unfold across a clade’s history, whether these changes influence diversification rates, and how morphological evolution tracks or lags behind ecological diversification. Australia offers an exceptional setting for studying such dynamics on a continental scale: its large size, long period of isolation, and dramatic climatic transitions—from rainforests to increasingly arid environments—have created a unique stage for evolutionary radiations (Byrne et al. 2008; Byrne et al. 2011; Pepper and Keogh 2021). The transition from widespread, relatively stable mesic environments during the early to mid-Cenozoic to the pronounced development of semi-arid and arid biomes starting approximately 15 Ma is hypothesized to have been a major driver of diversification in many Australian faunal groups (Byrne et al. 2011; Pepper and Keogh 2021). This time period is characterized by the expansion of the arid zone and fragmentation of mesic habitats (Byrne et al. 2008; Byrne et al. 2011), with mesic-adapted lineages either going extinct, persisting in climatically stable refugia (e.g., rocky escarpments), or invading the arid zone (Crisp et al. 2004; Fujita et al. 2010; Moritz et al. 2016).

Squamate reptiles (lizards and snakes) are the most species-rich terrestrial vertebrate group in Australia with over a thousand described species inhabiting all biomes (Powney et al. 2010; Uetz et al. 2021). Among the several iconic and remarkably diverse radiations included in this group—including agamid lizards, elapid snakes, varanid lizards, skinks, and pythons—are the endemic Gondwanan Pygopodoidea geckos, comprising three families (Carphodactylidae, Diplodactylidae, and Pygopodidae). With a crown diversification age between 50–70 Ma, the Pygopodoidea pre-date the complete separation of Australia from Antarctica (∼35 Ma), marking them as ancient Gondwanan endemics (Brennan and Oliver 2017; Kennett et al. 2018; Skipwith et al. 2019; Skipwith and Oliver 2023). In contrast, the remaining ∼700 immigrant lineages arrived from Asia and Melanesia between the Oligocene and Miocene, each with mesic-adapted ancestors and crown ages between 10–33 Ma (Oliver and Hugall 2017). The discrepancy between old stems and young crowns implies that much of the earlier Pygopodoidea diversity was lost, consistent with a mass turnover of ancestrally mesic-adapted lineages around the Eocene-Oligocene boundary (∼34 Ma)—a period of rapid global cooling and widespread contraction of rainforest habitats across Australia (Martin 2006; Hutchinson et al. 2021). Following this period, Pygopodoidea diversity recovered in conjunction with the arrival of immigrant squamate lineages, suggesting that habitat restructuring provided ecological space that was filled by both recovering endemic and new lineages that were dispersing into Australia (Brennan and Oliver 2017; Oliver and Hugall 2017).

In this study, we focus on the Diplodactylidae, the most species rich family of the Pygopodoidea. Diplodactylidae offers a great model system to study macroevolution within a regional gecko radiation. The family is distributed across Australasia, having expanded from Australia via two independent dispersal events to New Caledonia (NC) and Aotearoa New Zealand (ANZ) (Nielsen et al. 2011; Skipwith et al. 2016; Skipwith et al. 2019). The family comprises ∼200 described species, the majority of which are restricted to Australia (111 spp) followed by NC (66 spp) and ANZ (25 spp) (Uetz et al. 2021). A recent phylogenomic study of the group including ∼4200 ultraconserved element (UCE) loci confirmed that NC and ANZ each form a clade while the diversity in Australia comprises three clades: *Pseudothedactylus*, *Crenadactylus*, and the “core” Australian diplodactylids (Skipwith et al. 2019).

Members of Diplodactylidae occupy a wide range of habitats, from dry deserts and open spinifex (*Triodia*) grasslands, to wet tropical forests (Kluge 1967; Pepper et al. 2011; Oliver et al. 2012). Correspondingly, diplodactylids exhibit a remarkable breadth of ecomorphs (Sanders et al. 2008; Garcia-Porta and Ord 2013; Skipwith and Oliver 2023). Species on islands have independently evolved secondary diurnality—*Naultinus* in ANZ and *Eurydactylodes* in NC—and gigantism, including the largest extant gecko—*Rhacodactylus leachianus* with a maximum snout-vent length (SVL) of 255 mm—and an even larger extinct relative, *Gigarcanum delcourti* with an SVL of 370 mm (Heinicke et al. 2023). The Australian clades also display extraordinary morphological and ecological disparity, encompassing large predatory arboreal and cave species, saxicolous taxa that inhabit rocks and boulder fields, small ground-dwelling pointy-snouted termite specialists, and long-bodied spinifex specialists (Geneva 2015; Laver et al. 2017; Norris et al. 2021).

Variation in functional traits like body size, body shape, limb proportions, and foot morphology have been a great focus of study due to their incredible diversity in diplodactylids (Skipwith and Oliver 2023; Zimin et al. 2025; Pillai et al. 2025; Skipwith et al. 2026). However, tail morphology, a key functional trait among geckos and squamates that has been shown to serve a diverse array of functions—including balance, resource storage, communication, camouflage, and active defense—has been left out (Ramm et al. 2020; Griffing et al. 2021; Brennan et al. 2024, 2025). Diplodactylids exhibit some of the most extreme variation in tail morphology in any vertebrate family (Greer 1989; Skipwith et al. 2019; Green et al. 2024), yet no study has tested for ecological correlates with tail morphology across the group.

Here, we build a comprehensive phylogenomic dataset using anchored hybrid enrichment (AHE) loci, nuclear legacy exons (NLE), and ultraconserved elements (UCEs) and use this to generate a diplodactylid species tree to estimate the ancestral biome of diplodactylids based on current distributions. We then collect ecological and morphometric data including body and tail traits, and investigate the correlations between morphology and ecology. We test three hypotheses (Table 1): First, that the ancestral diplodactylid was arboreal and associated with mesic tropical biomes, with transitions into open, saxicolous, and spinifex-associated habitats tracking the expansion of open habitats across the continent. Second, that diversification rates differ among ecological states, with open-habitat and saxicolous lineages exhibiting higher diversification rates relative to arboreal lineages, consistent with increased ecological opportunity and habitat heterogeneity associated with the expansion of terrestrial and rocky environments during late Cenozoic environmental change in Australia. Third, that morphological traits differ among ecological states. Because tails contribute to balance, stabilization, and climbing performance in arboreal lizards (Jusufi et al. 2008), we predict arboreal lineages exhibit longer tails than other ecological states, and that arboreal and saxicolous lineages exhibit larger body sizes than ground-dwelling lineages, consistent with patterns documented in other lizard groups (Bauer and Sadlier 2000; Meiri 2008).

**Table 1:** Hypotheses tested, predicted patterns, and corresponding analyses.

| Hypothesis | Predicted pattern | Key analysis |
| --- | --- | --- |
| H1: Arboreal mesic origin | (a) Crown diplodactylids were arboreal and originated in mesic-tropical environments;<br>(b) transitions into ground, rock, and spinifex habitats track the progressive expansion of open habitats during Cenozoic aridification | Ancestral state estimations; paleobiome reconstruction |
| H2: State-dependent diversification | Open-habitat and saxicolous lineages exhibit higher net diversification than arboreal lineages, reflecting ecological opportunity associated with expanding terrestrial and rocky environments | state-dependent speciation and extinction analyses (SSE) |
| H3: Ecomorphological association | Arboreal lineages exhibit relatively longer tails, and arboreal and saxicolous lineages exhibit larger body size relative to ground-dwelling lineages | Phylogenetic MANOVA and GLS |

## Material and Methods

### Taxonomic sampling and sequencing

We sequenced 190 individuals representing 108 of the 111 recognized species of Australian diplodactylids. Sample preparation and sequencing methods were described in Tiatragul et al. (2023a) (see Supplementary Methods for details). Our sequencing targeted 388 anchored hybrid enrichment (AHE) loci, 5,023 ultraconserved elements (UCEs) loci and 42 nuclear legacy exons (NLEs) from the Squamate Conserved Loci (SqCL) panel (Singhal et al. 2017). We expanded our Australian dataset with published sequences from New Caledonian (n = 31 spp.) and Aotearoa New Zealand (n = 18 spp.) diplodactylids (Skipwith et al. 2019; Title et al. 2024), matching all sequences to common SqCL targets to ensure orthology. After alignment trimming and outlier masking, we retained 5,453 loci from 276 samples representing 171 species and recognized diplodactylid lineages. From the 276-sample dataset, we subsampled one representative per species and recognized subspecies for divergence dating, and nine species for a genus-level species tree analysis (see Extended Data 01). We trimmed and masked outliers separately for each subsampled dataset.

### Phylogenetic inference

We applied two phylogenetic inference approaches: a weighted summary coalescent method to estimate the species tree from gene trees, and a full Bayesian multispecies coalescent (MSC) model on genus-level representatives to resolve higher-level relationships.

#### Summary coalescent method and discordance analyses

We estimated maximum likelihood gene trees for each SqCL locus with IQ-TREE v2.3.6 (Minh et al. 2020) and used them as input for weighted ASTRAL (wASTRAL) v1.22.3.7 (Zhang and Mirarab 2022). We quantified phylogenetic concordance and discordance using the concordance vector (Ψ) framework (Lanfear and Hahn 2024), incorporating gene concordance factors (gCF), site concordance factors (sCF), and quartet concordance factors (qCF; see Supplementary Methods for details).

#### Inferring the Australian diplodactylid species tree using an MSC model

The placement of *Hesperoedura* within the core Australian clade is uncertain (Oliver et al. 2012; Skipwith et al. 2019); as a potentially early-branching lineage confined to relictual mesic forests of southwestern Australia, its position carries direct implications for ancestral biome estimation. We therefore estimated an alternative species tree for the core Australian diplodactylids using a full Bayesian MSC model in BPP v.4.7.0 (Yang 2015; Flouri et al. 2018) on the genus-level dataset (4,458 loci; 9 taxa), with *Crenadactylus horni* as the outgroup (Oliver and Sanders 2009). We explored nine and prior combinations across nine starting trees (81 total runs; see Supplementary Methods), assessing convergence by comparing posterior topology distributions across runs. We excluded 20 runs that did not yield a dominant topology and combined the remaining 61 to identify the MAP tree and generate an MCC tree. A final fixed-topology analysis obtained posterior estimates of and (see Supplementary Methods).

#### Divergence time estimation

We estimated divergence times with MCMCTree (Yang 2007) on the fixed wASTRAL topology, using four partitioned AHE datasets (n = 388 loci) and one unpartitioned UCE dataset (n = 100 loci). We selected 100 UCEs via a gene-shopping approach (Smith et al. 2018) for clock-likeness and low topological discordance (see Supplementary Methods), retaining the full AHE dataset. We conducted two analyses: a genus-level analysis calibrated with six squamate fossils (Table S1), and a full species-level analysis using five fossil and 13 secondary calibrations derived from the genus-level posterior (Table S2).

### Occurrence data

We compiled occurrence records for Australian diplodactylids from the Atlas of Living Australia (ALA) using the R package galah v.2.1.2 (Westgate et al. 2025), supplemented with data from the primary literature. Records obtained from ALA were visually inspected and manually corrected to reflect current taxonomy (see Extended Data 02).

### Ecological categorization

We characterized ecological states using two complementary schemes. Under the traditional ecotype framework, species were assigned to one or more predominant microhabitat categories: ground, rock, tree, and spinifex. Unique combinations of these categories were treated as discrete ecological states, resulting in eight observed habitat states (e.g., tree, tree–rock, spinifex–ground). To capture structural variation independent of habitat identity, we also developed a substrate-orientation scheme describing the typical orientation of occupied surfaces: horizontal (ground), vertical (tree trunks and rock faces), and narrow (twigs, foliage, and spinifex). As above, unique combinations were treated as discrete states, yielding six orientation categories (e.g., vertical, horizontal–vertical, narrow–vertical). These two schemes allowed us to distinguish substrate identity from structural orientation while accommodating ecological breadth in species occupying multiple microhabitats.

### Morphological data

We compiled a dataset of 19 linear measurements of morphological traits from the literature and museum specimens (n = 1367; see Extended Data 03), capturing gross morphological variation across the head, body, limbs, and tail following Brennan et al. (2024) (Table S3). Only specimens with complete measurements and original (non-regenerated) tail were considered (n = 509). All measurements were taken to the nearest 0.01 mm using a digital caliper (ABS Coolant Proof Caliper, Mitutoyo, Japan). To account for differences in overall body size, each trait was converted to a log shape ratio by dividing the raw measurement by the geometric mean of all measurements for that individual and then log-transforming the value. This transformation allows comparison of relative shape variation among species by removing the effect of size (allometry).

### Statistical analyses

#### Ancestral ecology estimation

We estimated ancestral ecological states (H1a) using discrete character evolution models in phytools v.2 (Revell 2024) in R v.4.5.2 (R Core Team 2025). For both habitat type and substrate orientation datasets, we compared equal-rates (ER), symmetrical (SYM), and all-rates-different (ARD) Mk models, as well as hidden-rates models (HRM) that allow unobserved rate heterogeneity among lineages (Beaulieu et al. 2013). To test whether ecological transitions follow constrained pathways, we also fitted an adjacency model in which transitions are permitted only between ecologically similar or intermediate states, reflecting the expectation that major ecological shifts occur stepwise rather than directly between strongly dissimilar states. We evaluated the fit of all models (i.e., ER, SYM, ARD, HRM, and adjacency model) to each dataset separately using AIC, and estimated marginal ancestral states under the best-supported model for each ecological dataset.

#### Paleobiome reconstruction and historical biogeography

To estimate the ancestral biome and biogeographic history of the clade (H1b) across major biomes on the Australian continent, we fitted a time-stratified dispersal-extinction-cladogenesis (DEC) model in the BioGeoBEARS v.1.1.3 (Matzke 2013) using the Köppen-Geiger biome classification scheme (Beck et al. 2018) to define relevant climatically-coherent ecoregions. This method is appropriate for the group compared to models that hierarchically define geographic regions and their constituent biomes (e.g., Landis et al. 2021), because each biome in Sahul is represented by a single contiguous area. We chose the DEC model over alternative range evolution models (e.g., DIVA and BayArea) as we were interested in the effect of range division during cladogenesis, but not in the effect of widespread sympatry, which is unlikely across the large spatial scale considered here. We estimated present-day biome occupancy per species by overlaying occurrence records with present-day biome maps (Beck et al. 2018); species were assigned to a biome if at least 10% of records fell within it, to a maximum of four biomes per species. Biomes have shifted their distribution dramatically across Australia during the period of diplodactylid diversification. To account for this, we defined time-stratified constraints on ecoregion occupancy and inter-regional dispersal at 10 Ma intervals from 0–50 Ma based on the connectivity of Köppen-Geiger ecoregions through time. We estimated historical location and connectivity of ecoregions from a paleoclimatic model of monthly mean temperature and precipitation at 1-degree resolution (Li et al. 2022), downscaled here to 0.1-degree resolution using the CHELSA algorithm (Karger et al. 2017). We also fitted a geography-only DEC model as a sensitivity check on biome classification (see Supplementary Methods for details).

#### State-dependent diversification modeling

We tested whether diversification rates differed among ecological states (H2) using state-dependent diversification models implemented in hisse 2.1.11 (Beaulieu and O’Meara 2016). We fitted models using the full time-calibrated phylogeny and complete ecological dataset to ensure accurate estimation of transition dynamics and hidden-state structure across the entire clade. We treated ecological states as the focal explanatory trait, and consolidated original habitat categories into three states: open terrestrial (ground- and spinifex-associated), arboreal (tree-dwelling), and saxicolous (rock-associated). We specified state-specific sampling fractions to account for incomplete taxon sampling (open terrestrial = 0.94; arboreal = 0.64; saxicolous = 0.94).

We fitted a suite of models including: (i) a “trivial null” MuSSE model assuming equal diversification rates across states, (ii) a full MuSSE model allowing state-dependent diversification but no variation within state, (iii) a character-dependent MuHiSSE model incorporating hidden rate traits, and (iv) character-independent diversification (CID) models with two to six hidden states, in which diversification rates are not linked to observed ecological states. The most complex CID6 matched the complexity of the MuHiSSE model with six diversification parameters. In all cases, we fixed the extinction fraction across observed and hidden states (Beaulieu and O’Meara 2016). For all models, we ran 40 maximum likelihood optimizations, each initiated from different starting values (Nakov et al. 2019). We assessed model fit using AIC.

We estimated marginal ancestral ecological states and diversification rates from all models that had >=1% Akaike weight using MarginReconMuHiSSE. We extracted model-averaged estimates of speciation (𝜆), extinction (𝜇), and net diversification (r) for both terminal tips and internal nodes. To avoid bias in ancestral state assignment caused by pruning nested non-Australian radiations, we estimated marginal ancestral ecological states on the full phylogeny. We excluded diversification rates and node assignments associated with NC and ANZ clades to retain only Australian taxa for downstream analyses. We assigned internal nodes to ecological states when the highest marginal posterior probability exceeded 0.66 and excluded ambiguous nodes.

#### Morphological association with ecological state and phylogenetic signal

To characterize the major axes of morphological variation, we conducted a principal component analysis (PCA) on the full 19-trait dataset using the FactoMineR R package (v.2.11) to identify the major axes of morphological variation. We then retained the 12 highest-loading traits for further comparative analyses—balancing representation of morphological variation with multivariate model performance—spanning six functional modules for comparative analyses: tail (tail length, tail width), body (body width, pelvic height), forelimb (upper and lower arm length), hindlimb (upper and lower leg length), head (head depth, posterior skull length, neck length), and geometric mean of size.

To test whether ecological state explains morphological variation (H3), we fitted multivariate phylogenetic GLS models using mvgls in the R package mvMORPH v.1.2.1 (Clavel et al. 2015), with ecological state as a fixed effect and phylogenetic relatedness incorporated via Pagel’s. We evaluated statistical significance using permutation-based MANOVA (Pillai’s trace, 1000 permutations), with pairwise post-hoc comparisons using BH correction (Benjamini and Hochberg 1995). We additionally fitted univariate phylogenetic GLS models for size and tail length to test directional predictions (see Supplementary Methods for details).

We quantified multivariate phylogenetic signal using Blomberg’s 𝐾_mult_ (Blomberg et al. 2003; Adams 2014), with the arithmetic (𝐾_𝐴_) and geometric (𝐾_𝐺_) means of K eigenvalues computed to assess whether signal was uniformly distributed across trait dimensions (Mitteroecker et al. 2025).

## Results

### Phylogenetic relationships of Australian diplodactylids

#### Summary coalescent method

Our full wASTRAL analysis corroborated previous results that Australian genera do not form a monophyletic group: *Pseudothecadactylus* was recovered as sister to the NC diplodactylids, while *Crenadactylus* is sister to the clade containing both the ANZ diplodactylids and the core Australian diplodactylids (*Amalosia*, *Hesperoedura*, *Oedura*, *Strophurus*, *Diplodactylus*, *Lucasium*, and *Rhynchoedura*) (Figure 1 a; Fig. S1). Our results also show that relationships among the core Australian diplodactylids remain difficult to resolve. While we found that *Diplodactylus*, *Lucasium*, and *Rhynchoedura* are a well-supported clade, concordance vector analyses show that the gene tree and site concordance is low (gCF < 30; sCF < 50) at deeper branches, including the splits between *Pseudothecadactylus*–NC clade and between *Crenadactylus*–ANZ + core Australian clade, indicating high gene tree/site conflicts (Extended Data 04; Extended Data 05).

**Figure 1:**
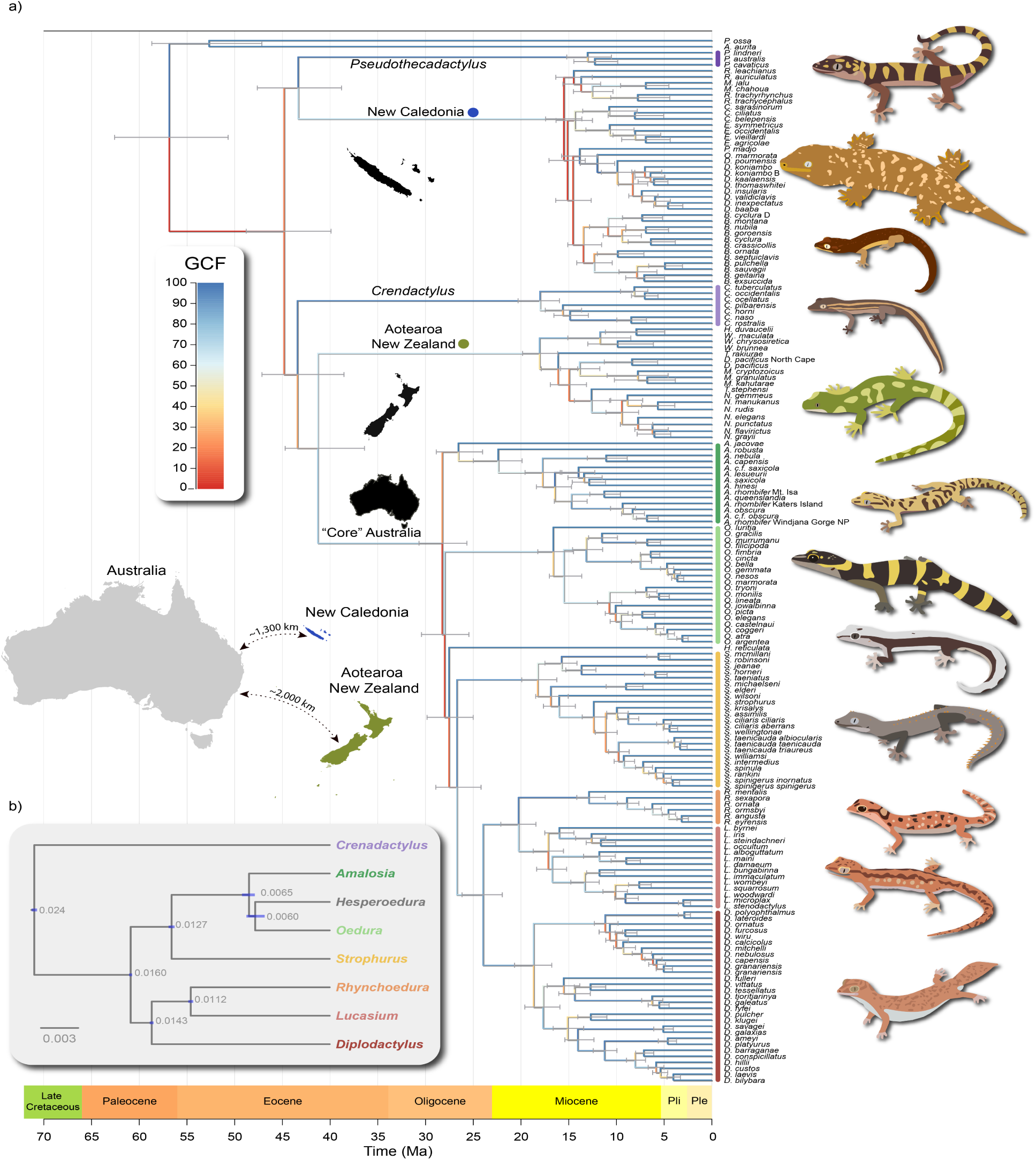
(A) Time-calibrated phylogeny of Australian diplodactylids and outgroup estimated in MCMCTree using Anchored Hybrid Enrichment (AHE) loci and ultraconserved element (UCE) loci based on the topology from summary coalescent method (wASTRAL). Node error bars represent 95% highest posterior density (HPD) intervals. Branch colors represent gene concordance factor. Inset maps show geographical separation between Australia, New Caledonia, and Aotearoa New Zealand. Color bars next to genera indicate Australian clades. (B) The maximum a posteriori tree from BPP shows a different topology. Node heights (𝜏) are the expected number of substitutions per site and are given at each node, with bars showing the respective 95% HPD intervals.

#### MSC method

Results from the BPP analyses on 4,428 loci dataset show a topology that is slightly different from the wASTRAL species tree (Figure 1 b). The MAP topology for all sets of priors is the same, with *Strophurus*, *Amalosia*, *Hesperoedura* and *Oedura* forming one clade and *Rhynchoedura*, *Lucasium*, and *Diplodactylus* forming the sister clade (Fig. S2; Table S4). Posterior values for the splits between *Amalosia* and *Hesperoedura* + *Oedura* clade, and for the split between *Hesperoedura* and *Oedura* mostly overlap. These results are consistent with rapid, successive divergences among these lineages, likely producing high levels of incomplete lineage sorting and leaving little phylogenetic signal to resolve their relationships (Degnan and Rosenberg 2009). However, mean values among *Rhynchoedura* and *Lucasium*, and between *Diplodactylus* and *Rhynchoedura* + *Lucasium* do not overlap, suggesting greater confidence in the divergence of these taxa.

### Divergence time estimation

The genus-level MCMCTree analysis placed the crown diversification of diplodactylids in the mid-Eocene (∼50 Ma; 95% HPD: 35–73 Ma), with the crown of the core Australian clade in the late Oligocene (∼30 Ma; 95% HPD: 20–42 Ma) (Fig. S3–4). The subsequent species-level analysis based on four AHE and one UCE partitions (Fig. S5) produced broadly consistent results, placing the crown of Diplodactylidae in the mid-Eocene (∼45 Ma; 95% HPD: 40–50 Ma) and the crown of the core Australian clade in the Oligocene (∼28 Ma; 95% HPD: 26–31 Ma) (Figure 1). Estimated divergence times between *Pseudothecadactylus* and New Caledonian diplodactylids, and between *Crenadactylus* and the Aotearoa New Zealand and core Australian diplodactylids, were both ∼43 Ma (95% HPD: 39–48 Ma). We estimated a crown age for the New Caledonian radiation at ∼15 Ma (95% HPD: 13–18 Ma) and the crown age for ANZ radiation at 18 Ma (95% HPD: 16–20 Ma). Within the core Australian clade, crown divergences of individual genera were dated to the late Oligocene–Miocene (10–27 Ma). Compared to previous UCE-based estimates (Skipwith et al. 2019), which placed the crown of diplodactylids in the late Cretaceous (∼60 Ma; 95% HPD: 52–69 Ma), our results suggest a substantially younger age for both the crown of the family and the crown ages of individual genera across Australia, New Caledonia, and Aotearoa New Zealand.

### Ancestral ecology and substrate orientation

Among ecological transition models, the equal-rates (ER) model was strongly preferred (AIC = 256.14, AICw = 0.99) over SYM (AIC = 266.36), ARD (AIC = 272.15), and constrained adjacency models (AIC = 280.31), and models incorporating hidden rate classes were not supported (Table S5). Under the ER model, marginal ancestral reconstruction estimated the ancestral diplodactylid was >99% tree-dwelling (Figure 2 a). Similarly, the most recent common ancestor for the core Australian diplodactylids was also tree-dwelling (>99%).

**Figure 2:**
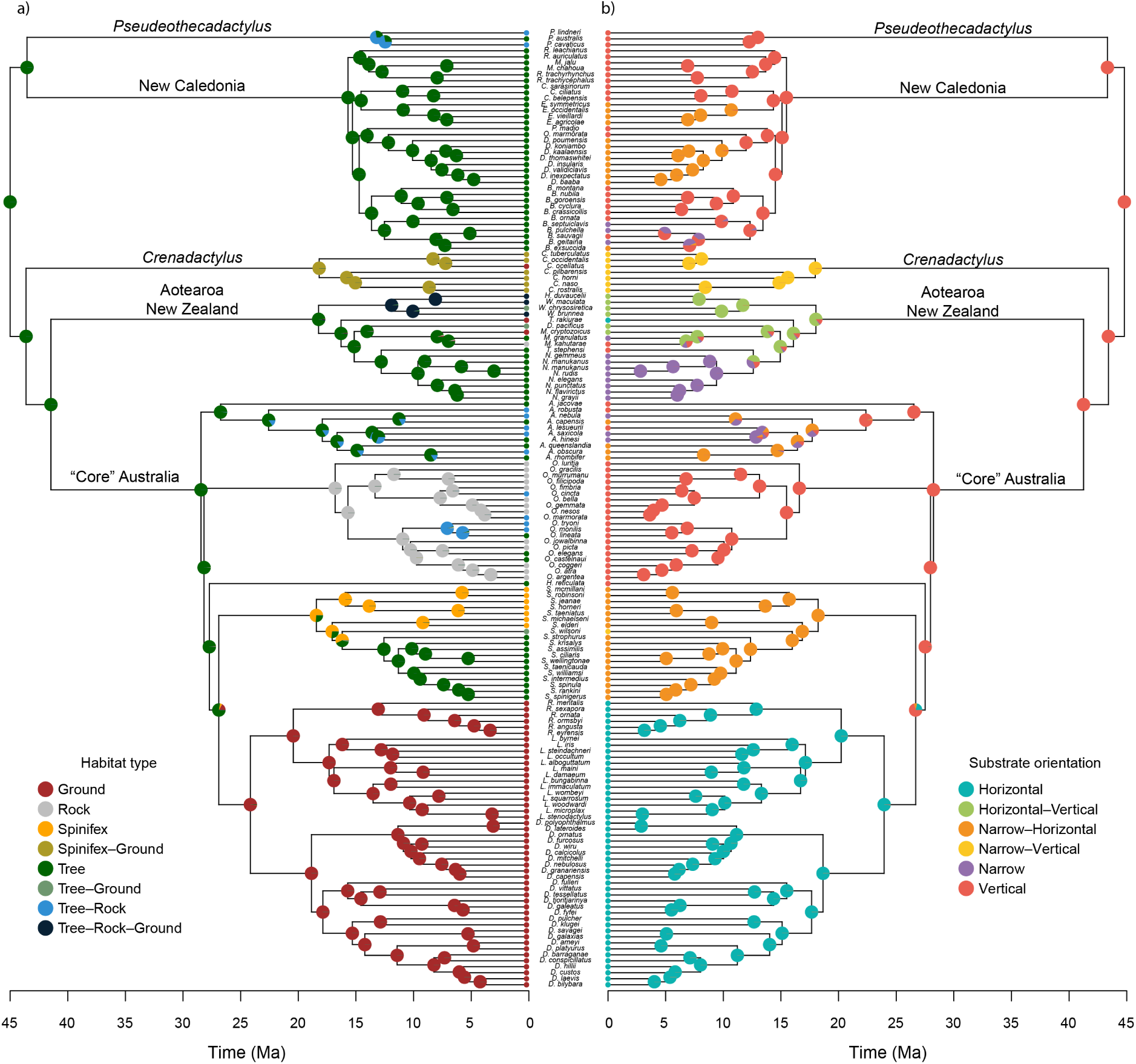
Ancestral state estimations of ecological states including (a) habitat type and (b) substrate orientation type. Pie charts at the nodes represent the probability of each state. Colors at tips represent present-day states.

For substrate orientation, model support was weaker, with ER (AIC = 203.09, AICw = 0.34), SYM (AIC = 201.99, AICw = 0.59), and ARD (AIC = 206.04, AICw = 0.08) models statistically indistinguishable (ΔAIC < 2), whereas the adjacency model performed poorly (AIC = 280.31; Table S6). Despite uncertainty in transition structure, ancestral state estimates were identical under ER and SYM models, consistently reconstructing a vertical substrate orientation for the ancestral nodes for crown diplodactylids and crown Australian clades with high confidence (Figure 2 b). Given this robustness, we reported results under the ER model. These ancestral state estimates indicate that early diplodactylids were arboreal and vertically oriented, with transitions into other ecologies and substrates occurring later in the clade’s history.

Australian diplodactylid clades showed greater ecological diversification than non-Australian lineages, with transitions into ground-, spinifex-, and rock-associated ecologies (e.g., *Crenadactylus*, *Diplodactylus*, *Rhynchoedura*, *Lucasium*). In contrast, New Caledonian taxa remained exclusively arboreal, while Aotearoa New Zealand lineages were mostly arboreal with limited ecological diversification, including habitat generalists (*Woodworthia*) and two predominantly ground-dwelling species (Figure 2 a). We found a similar pattern for substrate orientation: Australian clades showed greater diversification, including repeated transitions onto horizontal and narrow substrates and their combinations. New Caledonian species occupied primarily vertical, narrow, and narrow–vertical substrates, while Aotearoa New Zealand clades largely used mixed substrate orientations, most commonly narrow–vertical and horizontal–vertical combinations (Figure 2 b).

#### Ancestral biome

In the present-day, diplodactylid taxa are found across 11 different Koppen-Geiger ecoregions, though predominantly in semi-arid (BSh and BSk), arid desert (BWh and BWk) and tropical savanna biomes (Aw). The DEC model with time-stratified constraints on biome occupancy and dispersal from paleobiome reconstructions (Fig. S6), shows that range evolution in diplodactylids was characterized by low rates of dispersal among biomes (d = 0.02) and slightly higher rates of extirpation in biomes (e = 0.027), with dispersal attenuated by geographic distance (x = −0.16) and strongly influenced by climatic similarity (n = −0.44). Sympatric speciation, defined for range evolution models in the broad sense of speciation within the same region (Matzke 2013) and conceptually consistent with other modes of speciation within a single region (e.g., vicariance), accounts for the majority of speciation events. Narrow sympatry, defined in DEC models as when a species occupying a single biome diverges into two species in the same biome, accounted for 50% of cladogenetic events. Roughly 40% of cladogenetic events involved subset sympatry, in which a widespread ancestor diverged into two species, one within the complete range of the parent and one with only a single ecoregion of the parent. Whereas vicariant speciation across ecoregions accounted for only 11% of cladogenetic events.

The model did not find strong support for a single ancestral state, yet states that contained the tropical savanna biome were given higher probabilities (Figure 3; Extended Data 06). Similarly, biome states for many of the earliest divergences in the clade were highly uncertain, with support being shared among different possible state combinations. However, from the Miocene onward, we found less ambiguous support for biome states. Across 50 biogeographic stochastic maps from the DEC model, we find strongest support for a history of early occupancy of the tropical savanna biome, before spreading into semi-arid biomes in the Oligocene, and colonization of arid desert biomes (BWh and Bwk) and other temperate biomes in the Miocene by the clade comprising *Rhynchoedura*, *Diplodactylus*, and *Lucasium*. The exact timing of this process is not well resolved. Overall, across stochastic maps we found an average of 202.9 ± 12.4 transitions between all biome states, with the most common transition being the expansion from semi-arid to semi-arid + arid desert (12%), with 37.1% of all biogeographic events being dispersal to arid deserts from another biome.

**Figure 3:**
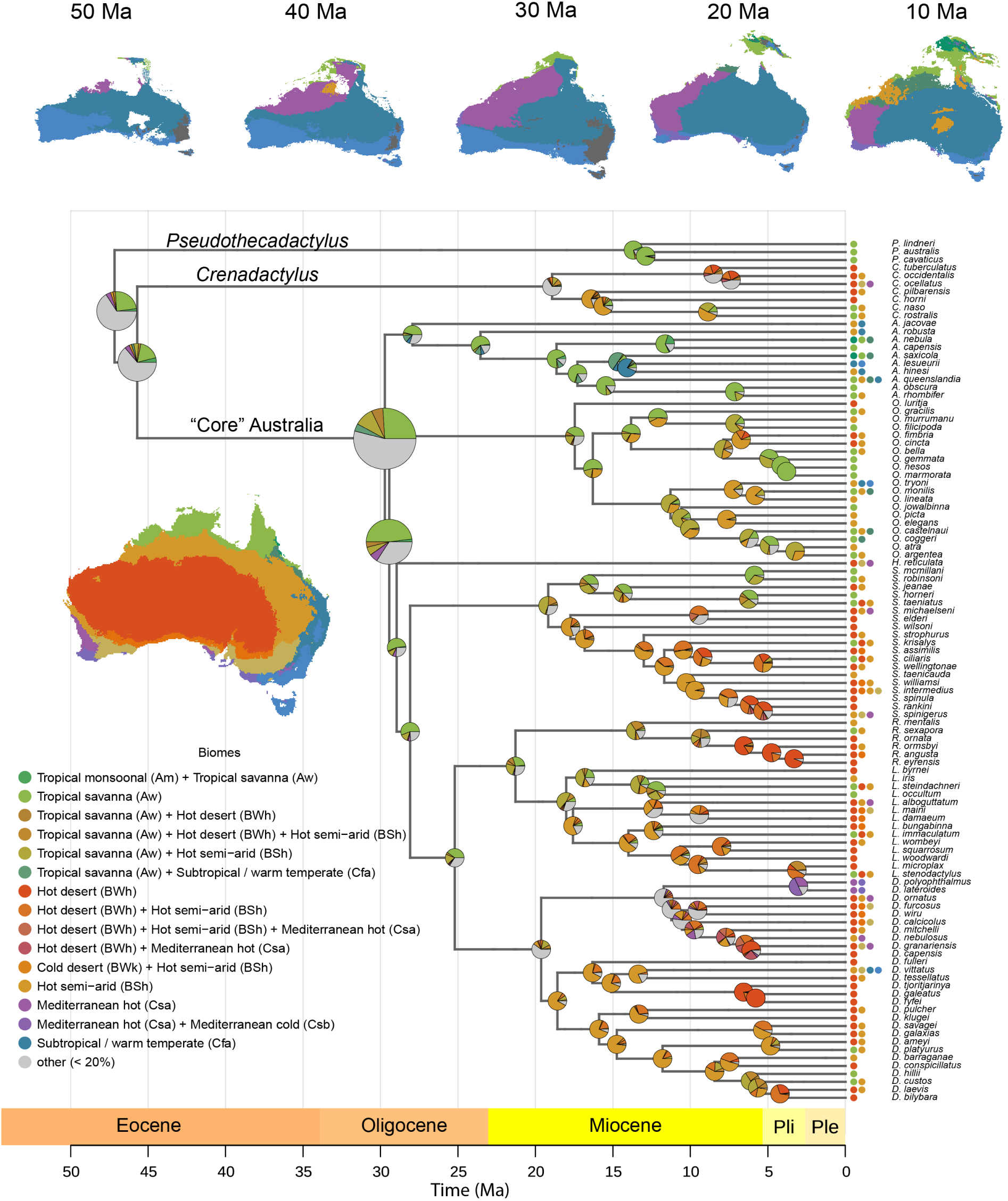
Ancestral biome estimates for Australian Diplodactylidae geckos according to the DEC model. Pie charts at the nodes represent probability from 50 stochastic maps. Only states or combinations of states that represent at least 20% of the nodes are shown. Colored circles at tips represent present-day state(s). Size of pie charts in ancestral nodes enlarged for readability.

The four-state geography model (northern, central, southeast, southwest) mirrors the 11-state biome model. Those results show similar uncertainty at deep nodes in the phylogeny; however, favor states that contain northern or central regions (Fig. S7). This is similar to support for the tropical and semi-arid biomes in the 11-state model at deep nodes, which are more widespread in the north and central parts of Australia, both in the present-day and historically (Fig. S8).

### State-dependent diversification and ecological effects

Model comparison supported character-independent diversification (CID) as the best explanation for rate heterogeneity across Diplodactylidae. CID3, with three hidden states, was the best-supported model (AIC = 1178.3, AICw = 0.42; Table S7), followed by CID4 (AIC = 1178.7, AICw = 0.35), CID5 (AIC = 1180.4, AICw = 0.15), CID6 (AIC = 1182.4, AICw = 0.05), and CID2 (AIC = 1183.7, AICw = 0.3). The character-dependent MuHiSSE model was not favored (ΔAIC = 22.6, AICw = 0), as were both MuSSE models (ΔAIC > 30). These results indicate that diversification rate heterogeneity exists within the radiation but is decoupled from observed ecological state: open, arboreal, and saxicolous lineages do not significantly differ in diversification rates after accounting for unmeasured hidden-states.

CID model-averaged marginal mean speciation rates per observed ecological state showed moderate differences between ecological states. Saxicolous lineages had slightly higher mean speciation rate (𝜆 = 0.140, 95% CI: 0.126–0.152), followed by arboreal lineages (𝜆 = 0.138, 95% CI: 0.125–0.151), whereas open-habitat lineages showed slightly lower rates (𝜆 = 0.115, 95% CI: 0.104–0.126; Table S8). Net diversification rates showed the same pattern with speciation as extinction rates were effectively zero across all states (Table S8), consistent with known difficulty in estimating extinction from extant-only phylogenies.

### Morphological evolution in diplodactylid geckos

Principal component analysis of the full 19-trait dataset revealed that the first three components explained 69% of total morphological variation (Figure 4 a; Fig. S9a). PC1 (30.5%) was dominated by tail width (37.3%), tail length (16.9%), and body size (16.6%). PC2 (25.8%) primarily reflected variation in body size (56.5%), with minor contributions from pelvic height (12.3%) and tail length (6.8%). PC3 (12.7%) was dominated by tail length (37.3%) and limb proportions, including lower arm (9.9%), upper arm (9.3%), and lower leg (7.4%). Across these axes, the largest contributors to morphological variation were body size, tail width, and tail length (Figure 4 b). These axes informed selection of the traits used in subsequent comparative analyses.

**Figure 4:**
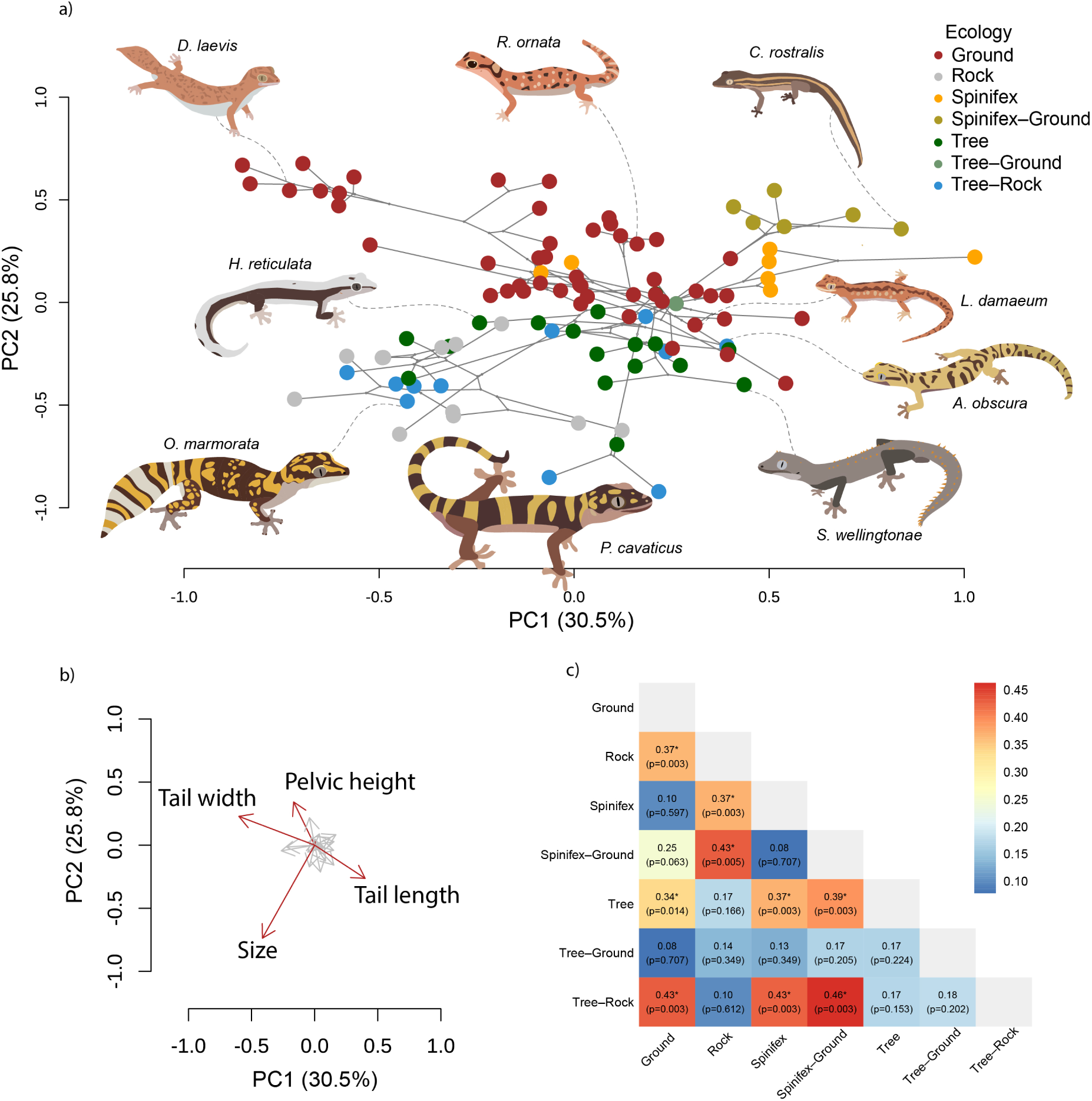
Morphological differentiation among ecological states in Australian diplodactylids. (A) Phylomorphospace of the first two principal components of morphological variation. Points represent species colored by ecological state, and connecting lines depict phylogenetic relationships. Axis labels indicate the proportion of variance explained by each component. (B) Principal component loadings showing trait contributions to the major axes of morphological variation. Vector length and direction indicate the magnitude and orientation of each trait’s influence in morphospace. (C) Pairwise phylogenetic MANOVA comparisons among ecological states based on Pillai’s trace. Cells show effect sizes (Pillai statistic) and Benjamini–Hochberg adjusted p-values; asterisks indicate significant differences. Color intensity reflects the magnitude of morphological differences between ecological states.

We found that morphological traits differed significantly among ecological states (Pillai’s trace = 1.11, p < 0.001). Pairwise comparisons showed the greatest difference between ground-associated (ground, spinifex, spinifex–ground) from non-ground lineages (rock, tree, tree–rock; Pillai’s trace 0.34–0.46, 𝑝_adj_ ≤ 0.007; Figure 4 c). Within these broad groupings, we found no statistically significant differences: ground-associated lineages (ground and spinifex) did not differ from one another (Pillai’s trace = 0.10, 𝑝_adj_ = 0.597), nor did non-ground lineages (rock, tree, tree–rock; Pillai’s trace 0.09–0.17, 𝑝_adj_ > 0.17). These results indicate that morphology varies with broad ecological states and is tightly linked to surface-use.

Morphological evolution was also strongly structured by phylogenetic relatedness. We detected multivariate phylogenetic signal across morphology (𝐾_mult_ = 0.81, p = 0.001). However, phylogenetic signal was unevenly distributed across trait dimensions, with the arithmetic mean exceeding the geometric mean (𝐾_𝐴_ = 0.61, 𝐾_𝐺_ = 0.42; 𝐾_𝐴_/𝐾_𝐺_ = 1.46), indicating that some axes of variation are more strongly phylogenetically structured than others (𝑍_𝐾mult_ = 10.73; 𝑍_𝐾𝐴_ = 6.97; 𝑍_𝐾𝐺_ = 10.94; Fig. S9b—c). Consistent with this pattern, the dominant axes of variation were driven by tail width and body size (PC1–PC2), with tail length and limb proportions contributing to the third axis (PC3). Together, these results indicate that morphological evolution is strongly structured by ancestry but exhibits trait-specific ecological divergence.

Univariate phylogenetic models showed trait-specific ecological effects. Size differed significantly among ecological states (df = 6, F = 7.93, 𝑝_adj_ < 0.0001). We report marginal means as back-transformed geometric mean size to mm, with contrasts expressed as size ratios relative to ground-dwelling lineages (6.41 mm). Rock-associated (9.40 mm; 1.46×, 𝑝_adj_ = 0.003), arboreal (8.95 mm; 1.39×, 𝑝_adj_ = 0.008), and tree–rock lineages (9.89 mm; 1.54×, 𝑝_adj_ < 0.001) were all significantly larger than ground-dwelling species, while spinifex–ground lineages were significantly smaller (4.67 mm; 0.73×, 𝑝_adj_ = 0.007; Table S9). In contrast, tail length did not differ significantly among ecological states (df = 6, F = 1.89, 𝑝_adj_ = 0.09; Table S10). These results indicate that morphological differentiation among ecological states is driven primarily by size, while tail length is phylogenetically conserved.

## Discussion

Australian diplodactylid geckos represent one of the most species-rich, ecologically diverse, and oldest reptile radiations on the continent, yet the processes that generated this diversity have remained poorly understood. By integrating phylogenomics, ancestral state estimation, diversification modeling, and comparative morphology, our analyses provide a comprehensive macroevolutionary framework linking environmental change, ecological expansion, and morphological diversification across deep time. Three primary conclusions emerged. First, (H1) Australian diplodactylids likely originated in mesic, arboreal environments and subsequently underwent repeated ecological transitions as the continent became drier and open habitats became available. Second, (H2) diversification rates exist across the radiation but are independent of observed ecological states, with character-independent models favored over character-dependent models. Third, (H3) morphological evolution reflects a combination of strong phylogenetic structure and ecological divergence, with body size varying among ecological states while tail morphology remains comparatively conserved. Together, these results support a model in which continental environmental change was associated with ecological opportunity that influenced diversification through habitat transitions and selective morphological differentiation.

### Environmental change and the temporal assembly of the radiation

Our divergence estimates indicate that the crown group of Diplodactylidae originated in the early Eocene (∼45–50 Ma), when Australia remained connected to Antarctica and South America as part of the southern Gondwanan landmass. At this time, much of the region supported temperate to subtropical forest ecosystems (Martin 2006). Over the next 45 Ma Australia migrated faster than any other continent, travelling more than 3000 km towards the equator (Quigley et al. 2010). As Australia drifted rapidly northward, the continent experienced profound climatic shifts that reshaped terrestrial ecosystems—from warm, humid Eocene rainforests to cooler and increasingly seasonal Oligocene environments (Martin 2006; Amoo et al. 2022), and eventually the widespread expansion of open woodlands, shrublands, and desert biomes during the Miocene (Byrne et al. 2008; Pepper and Keogh 2021).

Our inferred timing of diversification within Australian diplodactylids aligns with these environmental transitions. Core Australian diplodactylids originated in the Oligocene (∼28–30 Ma), roughly 10 Ma later than previous estimates (Skipwith et al. 2016; Skipwith et al. 2019). This timing broadly corresponds to a period of accelerated climatic and biotic changes that occurred during and immediately after the transition between the Eocene and the Oligocene (Brennan and Oliver 2017; Nge et al. 2020; Hutchinson et al. 2021). Diversification of extant groups occurred much later during the Miocene. The earliest diversification within the Australian radiation occurs during the late Oligocene ∼27 Ma in *Amalosia*, a genus restricted today to the wetter periphery of the continent.

By contrast, diversification within the arid-adapted genera (*Oedura*, *Rhynchoedura*, *Strophurus*, *Diplodactylus*, and *Lucasium*) occurred later, with most diversification occurring ∼15 Ma in line with the first signs of widespread continental aridity (Byrne et al. 2008; Pepper and Keogh 2021). This timing strongly suggests that the tempo of diplodactylid evolution is associated with this ecological transition and the eventual emergence of the arid biome. Rather than representing a single burst of diversification, the radiation appears to have assembled through successive phases of lineage expansion linked to the emergence and spread of new ecological opportunities across the continent. Several other Australian radiations share this pattern of an Oligocene origin followed by Miocene diversification concurrent with aridification including reptiles (Rabosky et al. 2007; Pavón-Vázquez et al. 2022; Torkkola et al. 2026), insects (Owen et al. 2017; Heimburger et al. 2022; Li et al. 2024), and plants (Jabaily et al. 2014; Hammer et al. 2021; Hua et al. 2022), suggesting that continental aridification drove diversification across taxonomically unrelated groups.

Our estimates are younger than those in Skipwith et al. (2019) primarily because we excluded a secondary calibration prior on the pygopodoid crown and used partitioned AHE loci as well as UCEs chosen for clock-likeness and low topological discordance rather than taxon completeness alone; these differences affect rate estimates and consequently node ages. Topological differences between studies concentrate at the deepest backbone nodes, consistent with the low gCF and sCF values observed there (see Supplementary Discussion).

Biogeographic patterns outside Australia reinforce the importance of post-Eocene environmental change. Our phylogenetic reconstruction corroborates previous findings that the NC and ANZ diplodactylid radiations are far too recent to reflect the Cretaceous rifting of Zealandia from Australia (Mortimer et al. 2017). Instead, these clades almost certainly diverged from their Australian ancestors while still on the Australian continent and subsequently colonized the islands via long-distance overwater dispersal (Wallis and Jorge 2018; Crisp et al. 2019; Skipwith et al. 2019). Crown ages of 16–20 Ma follow the Oligocene Marine Transgression (also referred to as the Oligocene “Drowning” ∼23–25 Ma), when Zealandia experienced maximum inundation and terrestrial habitats were most restricted (Strogen et al. 2009; Wallis and Jorge 2018). Geological reconstructions indicate that substantial land re-emerged during the earliest Miocene (∼23–21 Ma), creating extensive terrestrial environments for the first time since peak drowning of Zealandia (Strogen et al. 2009). The close temporal correspondence between land re-emergence and diversification strongly supports a Miocene colonization scenario rather than ancient vicariance (Wallis and Jorge 2018). Together, these patterns highlight how regional geological and climatic dynamics have repeatedly structured opportunities for dispersal and diversification across the diplodactylid lineage.

### Ancestral ecology (H1a)

Ancestral state estimation indicates early diplodactylids were arboreal and associated with vertical substrates, consistent with the predominance of forested environments across Australia during the Eocene (Byrne et al. 2011; Pepper and Keogh 2021). From this ancestral condition, lineages repeatedly transitioned into alternative ecological states, including ground-associated, rock-dwelling, and spinifex-specialized habitats. The best-supported transition model indicates broadly equivalent transition rates among ecological states, suggesting that ecological diversification proceeded through repeated habitat shifts rather than strongly constrained evolutionary pathways.

The timing of these transitions closely mirrors the progressive expansion of open habitats and diversification of xeromorphic vegetation across Australia during the late Oligocene and Miocene (Byrne et al. 2008). As forests contracted and structurally heterogeneous environments spread, new ecological niches emerged for terrestrial and shrub-dwelling lineages. Independent origins of spinifex specialization in *Strophurus* and *Crenadactylus* further illustrate how environmental restructuring provided opportunities for ecological novelty (Crisp and Cook 2013; Doughty et al. 2016; Mulder et al. 2022). The diversification of spinifex-specialized lineages within both genera broadly coincides with the late Miocene diversification of *Triodia* itself (crown 7.0–8.8 Ma; Anderson et al. (2019)). These findings indicate that ecological diversification in Australian diplodactylids reflects dynamic habitat tracking and expansion in response to long-term continental environmental change rather than simple retention of ancestral ecological conditions.

Diplodactylids in NC and ANZ provide an important comparison. Lineages on these islands have remained predominantly arboreal but have independently evolved traits not observed in Australian diplodactylids, including gigantism, viviparity, and diurnality (Bauer and Sadlier 2000; Nielsen et al. 2011; Skipwith et al. 2016; Skipwith and Oliver 2023; Heinicke et al. 2023). This contrast may reflect differences in ecological context between island and continental systems. In NC and ANZ, the forest-dominated, predator-free environments may have facilitated the evolution of large-bodied, diurnal, or cold-adapted lineages (Skipwith et al. 2026), whereas the Australian continent hosts a diverse assemblage of predators and ecologically similar squamate lineages that may constrain the evolution of such traits. Instead, ecological diversification within the Australian radiation appears linked primarily to repeated transitions into terrestrial habitats as open habitats expanded across the continent. Ground-specialist diplodactylids are largely absent outside of Australia, reinforcing the idea that evolution of terrestrial morphotypes is linked to the availability of open arid habitats that characterize the Australian continent. Australian agamid lizards show a comparable pattern where several lineages, including the thorny devil (*Moloch*) and earless dragons (*Tympanocryptis*), are fully terrestrial in Australia (Brennan et al. 2025), yet none of their closest New Guinean relatives occupy terrestrial niches (Tallowin et al. 2019). Another example is in *Anolis* lizards where ground-dwelling species are common on the mainland but rare on islands (Huie et al. 2021). That open-habitat specialists arise repeatedly on the continent but not on forested islands, across unrelated clades, suggests niche availability contributes to ecological diversification more than clade identity.

### Ancestral biome (H1b)

Biogeographic reconstruction provides a complementary perspective on how environmental change structured the spatial and ecological assembly of the clade. The biogeographic range evolution model identified biomes that contained the tropical savanna of northern Australia as the most likely ancestral biome for Australian diplodactylids, followed by expansion into semi-arid and ultimately fully arid biomes. This pattern is consistent with our prediction that diplodactylids originated in relatively mesic environments and expanded into progressively drier environments tracking the geographic expansion of open and increasingly dry habitats through time (Byrne et al. 2011). Notably, modern Australian diplodactylids are largely absent from rainforest environments except for a small number of species in *Pseudothecadactylus* and *Amalosia*, whereas rainforest systems in NC and ANZ retain predominantly arboreal diplodactylid assemblages. In Australia, rainforest gecko communities are instead dominated by the closely related Carphodactylidae (Brennan and Oliver 2017), suggesting long-term ecological partitioning among Pygopodoidea lineages across biome types.

The inferred history of biome occupancy is highly dynamic, with an average of 202 transitions across the phylogeny, with over a third of these into desert environments. A major shift into arid environments was inferred for the *Rhynchoedura–Diplodactylus–Lucasium* clade near the Miocene onset, coinciding with continental drying — a pattern paralleled in agamid lizards, Pygopodoidea, and typhlopid snakes (Brennan and Oliver 2017; Tiatragul et al. 2023b; Brennan et al. 2025). Because biogeographic models are estimated from extant lineages, extinction of intermediate-biome lineages would further inflate inferred transition counts, reinforcing rather than contradicting the pattern of dynamic biome occupancy.

Despite clear directional expansion into arid landscapes, many arid-zone lineages trace their origins to ancestors that predate the modern arid zone (Pepper et al. 2006; Skeels et al. 2025). The clade comprising *Strophurus*, *Diplodactylus*, *Lucasium*, and *Rhynchoedura* dates to the Oligocene, but Australia’s arid interior developed progressively through the Miocene and Pliocene with some of the driest systems emerging as recently as the late Pleistocene ∼1 Ma (Pepper and Keogh 2021). Our paleoenvironmental reconstructions suggest that semi-arid climates persisted across northern Australia throughout the Cenozoic, potentially pre-adapting early diplodactylid lineages to subsequent aridification. However, very little climate proxy information is available for northern Australia during the early Cenozoic, and as such the degree to which arid biomes existed, and were functionally and structurally similar to arid biomes today, remains unknown. The geography-only model showed the most likely location for ancestral diplodactylids was in northern and central parts of Australia, pre-desertification of the continent. This suggests that microhabitat affinities not captured by broad scale climatic classifications may be important in defining species distributions over deep time scales. Nonetheless, the temporal mismatch between lineage origin and biome formation thus likely reflects progressive tracking of shifting climatic boundaries over time in a clade that is particularly well suited to persisting in dry conditions, rather than instability in biome association (see Supplementary Material for extended discussion).

### Ecological states and diversification dynamics (H2)

State-dependent diversification analyses indicate that rate heterogeneity exists within the Diplodactylidae radiation, but character-independent models outperformed character-dependent MuHiSSE, indicating that unmeasured factors explain rate variation better than our broad ecological classification. The three-state classification likely captures a coarser ecological axis than the factors actually driving speciation and extinction dynamics.

Rate heterogeneity may track geographic or microhabitat variation not captured by the broad ecological categories. For example, spinifex-specialist lineages vs. burrow-associated species within the open category, or rocky outcrop endemics vs. generalist saxicolous species. Alternatively, the three-state categorization may lump ecologically distinct lineages whose rate differences cancel out across states. The support for CID3 specifically (three hidden states) suggests a three-way rate structure in the radiation, but the biological correlates of these hidden states remain unidentified and represent a productive avenue for further investigation.

Several studies have documented mid- to late-Miocene turnover and show that biome change and aridification have been associated with both lineage turnover and episodic radiations in Australian taxa, including diplodactylids (Crisp and Cook 2013; Brennan and Oliver 2017; Owen et al. 2017; Tiatragul et al. 2023b). These paleoenvironmental changes are a plausible source of ecological opportunity that influenced diversification in diplodactylids by creating new ecological space and by fragmenting formerly continuous ranges (Brennan and Oliver 2017). Arboreal environments can promote niche partitioning and fine-scale microhabitat segregation, providing opportunities for sympatric divergence (Westeen et al. 2025). Although ecological transitions are well supported by ancestral state reconstruction, these shifts did not translate into detectable differences in diversification rates under the character-independent framework favored here. Saxicolous systems offer another mechanism. Rocky ranges are spatially discontinuous and harbor short-range endemic diplodactylid species, particularly scansorial taxa in *Pseudothecadactylus* and *Oedura* (Hoskin and Higgie 2008; Oliver et al. 2014; Laver et al. 2018). For other gekkonids, rocky outcrops and karst limestones generate similarly high levels of endemism through isolation (Grismer et al. 2018; Oliver et al. 2019). These environments can also function as climatic refugia during environmental fluctuations, facilitating divergence during periods of range contraction and expansion (Laver et al. 2018; Oliver et al. 2019). Open terrestrial habitats, including ground and spinifex systems, can foster diversification through substrate-specific adaptations (e.g., burrow use and spinifex association) not captured by our broad categorization, yet their broader geographic continuity may reduce opportunities for prolonged isolation relative to rock or vertically structured arboreal systems. This pattern reflects the landscape heterogeneity of large continental radiations and distinguishes them from island systems where ecological opportunity may be more constrained to a smaller set of habitats.

### Morphological evolution under phylogenetic constraint (H3)

Morphological analyses reveal significant but trait-specific differentiation among ecological states: multivariate phylogenetic models place species in distinct regions of morphospace corresponding to their major ecological category. However, strong phylogenetic signal in several traits indicates that ecological divergence is constrained by shared ancestry, as found in other Australian lizard radiations (Openshaw and Keogh 2014; Ashman et al. 2018; Tiatragul et al. 2024).

Ecological effects are not uniform across traits. Size differs significantly among ecological states, with several non-ground lineages exhibiting larger mean size relative to ground- and spinifex-associated taxa, a pattern repeatedly documented in reptile radiations (Huie et al. 2021; Riedel et al. 2024; Westeen et al. 2025; Oliver et al. 2026). Contrary to our prediction, we found no correlation between tail length and our broad ecological categories, despite striking variation in tail morphology across the family and the importance of tail length in overall trait space (PC1 and PC3).

These findings refine previous interpretations of morphological diversification within the family. Diplodactylids exhibit extraordinary diversity in tail morphology and function (Greer 1989), including defensive tail glands to shoot adhesive “goo” for defense, prehensile structures, and enlarged bulbous-shaped tails for fat storage (Bustard 1967; Nielsen et al. 2016; Green et al. 2024). Our results show that tail width dominates the PC1, while tail length contributes to PC3. Neither tail dimension shows a clear association with our broad ecological state. Instead, ecological differentiation appears to be expressed more strongly through variation in body size, suggesting that ecological divergence in Australian diplodactylids primarily involves changes in overall size rather than coordinated modification of limb or tail structures as documented in the radiation of *Anolis* lizards (Losos 2009; Huie et al. 2021). Together, these results show that ecological and phylogenetic processes both shape morphology in Australian diplodactylids, but their influence differs: ecology is correlated with overall body form and size, while tail morphology is constrained by phylogeny.

Future studies using 3D geometric morphometric approaches would provide a more complete characterization of tail shape and may reveal ecological associations undetectable using linear measurements alone (see Supplementary Discussion).

## Conclusion

The Australian diplodactylid radiation was assembled through the interaction of environmental change, ecological expansion, and phylogenetically structured morphological evolution. Long-term climatic transformation created new ecological opportunities; repeated habitat transitions contributed to diversification, though rate heterogeneity was largely independent of observed ecological state; and morphological adaptation occurred within constraints imposed by shared ancestry. Rather than a single adaptive burst, the radiation reflects prolonged ecological expansion across an evolving continental landscape.

This pattern highlights an important distinction between continental and island radiations. Within Diplodactylidae, the NC and ANZ clades provide a direct comparison: both remained predominantly arboreal, independently evolving traits such as gigantism, viviparity, and diurnality within forest-dominated environments, rather than undergoing the repeated habitat transitions that characterize the Australian radiation. Continental radiations may be shaped by the gradual emergence of ecological opportunity through environmental transformation rather than the rapid filling of discrete niche space (Yoder et al. 2010; Stroud and Losos 2016) — producing diversification through prolonged ecological expansion rather than competitive replacement. Australian diplodactylids exemplify this process: their history demonstrates that large continental radiations can unfold as extended, multi-phase responses to geographic and climatic change.

## Acknowledgements

We thank curators and collection managers from the South Australian Museum, Western Australian Museum, Queensland Museum, Australian National Wildlife Collection, Museum and Art Gallery of the Northern Territory, and Museums Victoria for providing access to specimens and many of the tissue samples used. ST thanks Adam Leache for help with BPP analyses, Robert Lanfear for help with concordance vector analyses, and Conrad Hoskin, Paul Doughty, Paul Oliver, Aaron Bauer and Wesley Read for helpful discussions about diplodactylids. We thank Siddhant Attavar for help with implementing new features in pipesnake and Elizabeth Broady for help with sequencing preparation. We thank and Australian BioCommons Leadership Share Program (ABLeS) for the providing computation and digital resources.

## Data Availability Statement

Associated data, script, and extended data are available on Dryad: 10.5061/dryad.jsxksn0r2 in the final manuscript.

