## Supplementary Material for "Aridification and habitat shifts are associated with diversification in Australian diplodactylid geckos"

Sarin Tiatragul[1,*], Ian G. Brennan[1,2], Alexander Skeels[1], Damien Esquerré[3], Stephen M. Zozaya[1], J. Scott Keogh[1], and Mitzy Pepper[1]

[1]Division of Ecology & Evolution, Research School of Biology, The Australian National University, Canberra, ACT 2601, Australia

[2]Biodiversity and Geosciences, Queensland Museum, PO Box 3300, South Brisbane BC, Queensland 4101, Australia

[3]Environmental Futures Research Centre, School of Science, University of Wollongong, Wollongong, NSW 2500, Australia

### Table of contents

Data availability 3

Extended Data 4

Supplementary Figures 5

Supplementary Tables 15

Supplementary Methods 26

References 37

#### Data availability

Associated data, script, and extended data are available on Dryad: 10.5061/dryad.jsxksn0r2

### Extended Data

Extended data is available online on Dryad and includes the following:

1. List of samples for phylogenomic analyses
2. Occurrence records
3. Raw morphological measurements of specimens
4. gCF/sCF/qCF annotated tree
5. Concordance vectors (gene, site, and quartet concordance factors)
6. BioGeoBEARS all-state node state probabilities

### Supplementary Figures

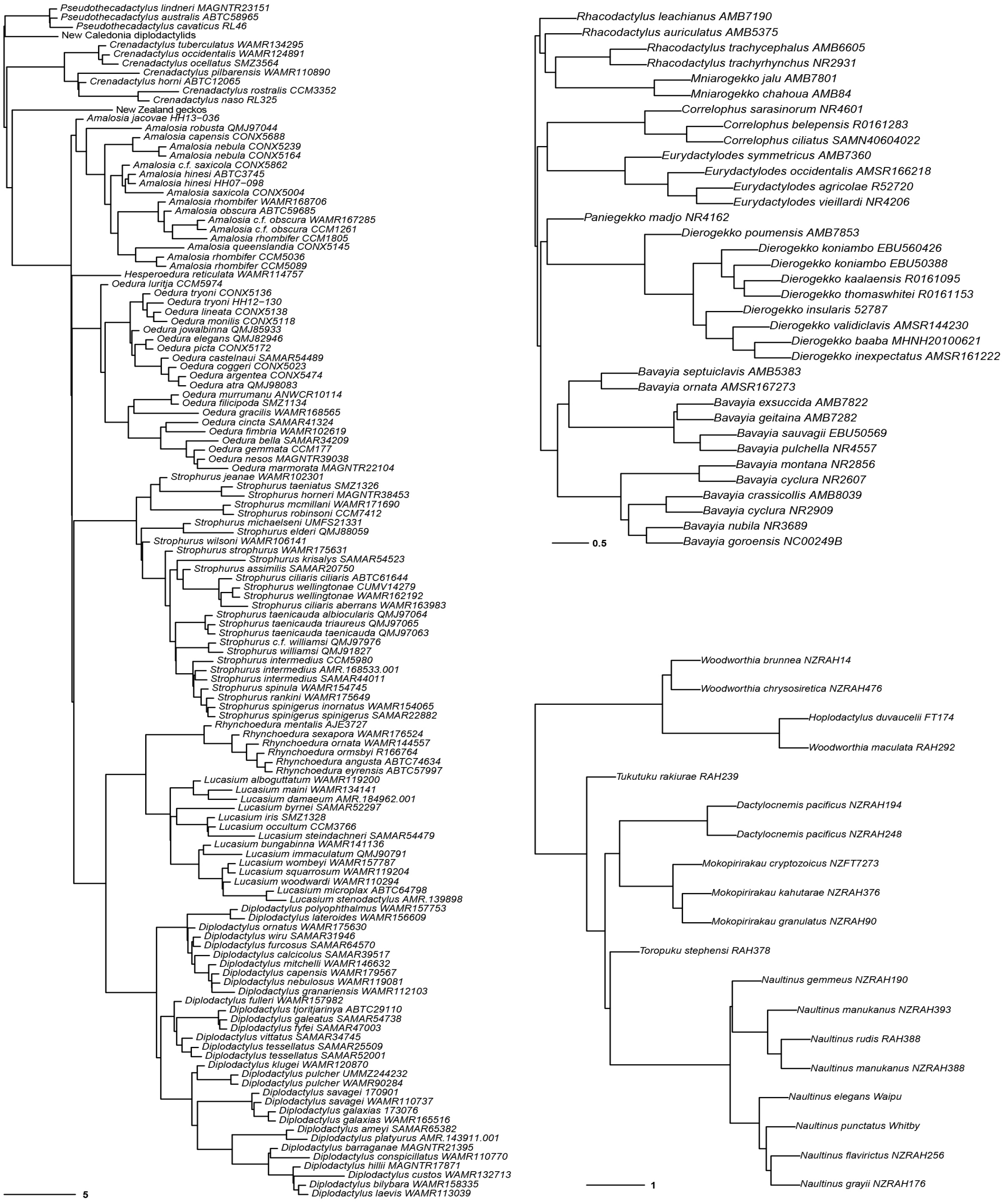

Figure S1. Phylogenomic tree inferred using summary coalescent method (wASTRAL) based on 5,391 SqCL loci maximum likelihood gene trees of a) a subset of Australian diplodactylids, b) NC diplodactylids, and c) ANZ diplodactylids. Scale bar shows branch length in coalescent units. Full trees are available in online repository.

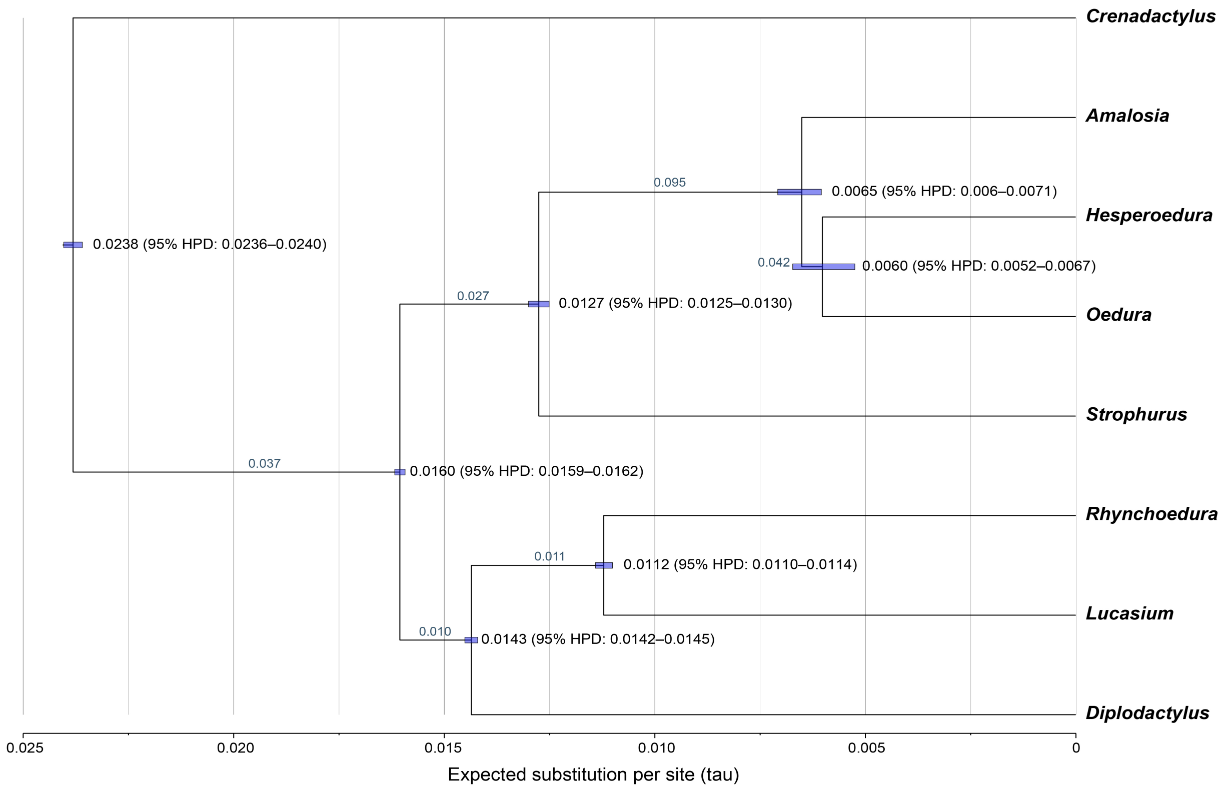

Figure S2. The maximum a posteriori (MAP) species tree for Australian core diplodactylids showing the parameter estimates (x 0.01). Node heights ($\tau$) are the expected number of substitutions per site and are given at each node, with bars showing the respective 95% HPD intervals. Posterior means of $\theta$ (population size) are shown along the branches. The prior used in the A00 analyses were $\theta \sim$ InvG(3, 0.1) for all populations and $\tau \sim$ InvG(3, 0.03). The tree is drawn with FIGTREE using the BPP output FigTree.tre

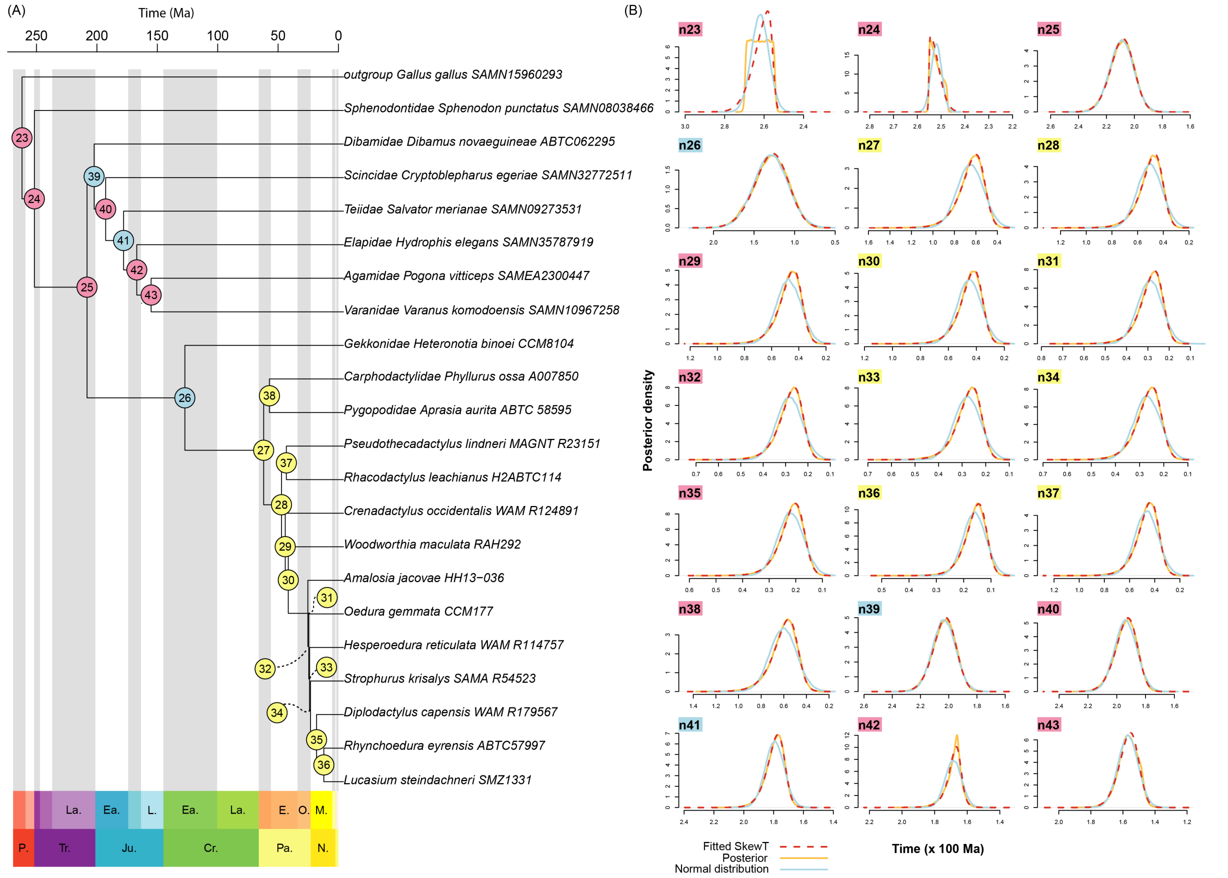

Figure S3. a) Calibration scheme for genus-level MCMCTree analysis. Fossil (pink) and secondary (yellow) calibrations are indicated at nodes. Blue nodes are uncalibrated. b) Posterior density (orange line) and fitted skew-T (dotted red line) distribution from each node. Blue normal distribution plotted as reference. Skew-T distributions are used as secondary priors for species-level MCMCTree analysis.

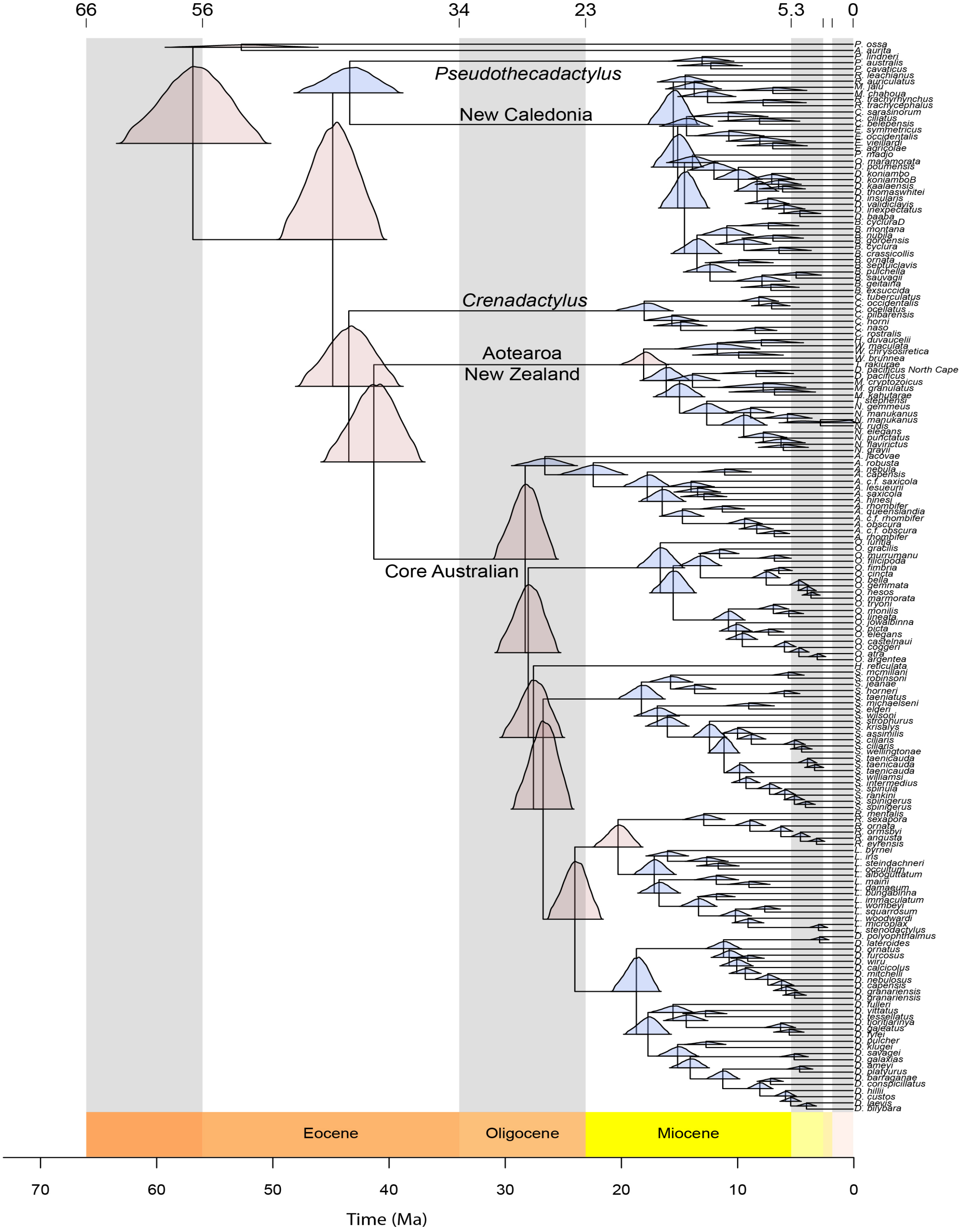

Figure S4. Results from MCMCTree with posterior density shown at the nodes. Pink nodes indicate secondary calibration with bounded priors. Blue nodes were estimated without calibration.

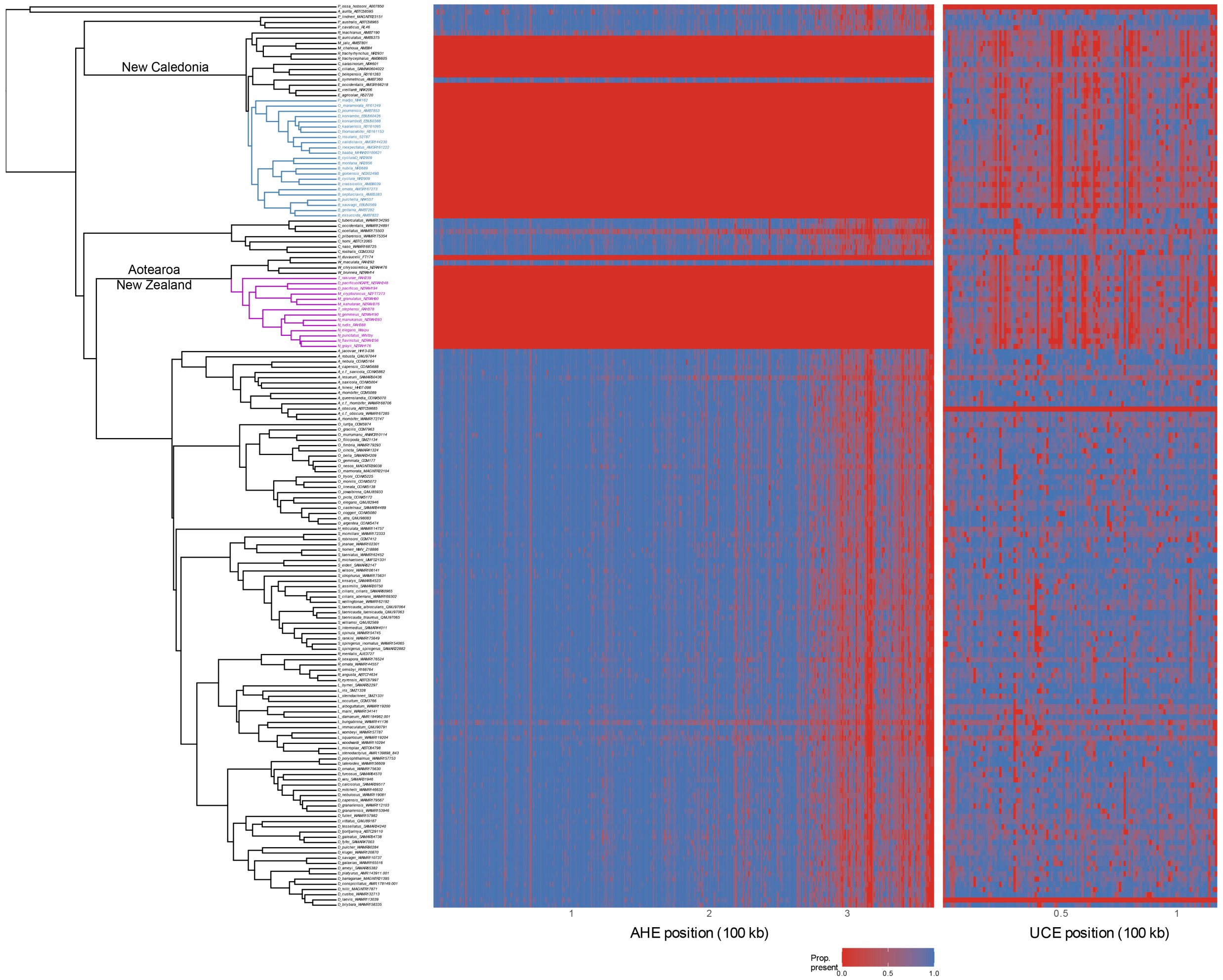

Figure S5. Alignment completeness across the 5-partitions MCMCTree analysis for 175 Diplodactylidae taxa. Left panel shows the phylogeny, and the two right panels show per-taxon sequence completeness as heatmaps for the combined AHE partitions 1-4 (middle) and UCE partition 5 (right), with taxa ordered to match the tree tip order. Cell color indicates the proportion of non-gap/non-missing sites within 1 kb bins (ranging from blue = high completeness to red = absent). Most New Caledonian and Aotearoa New Zealand taxa are represented only by UCEs.

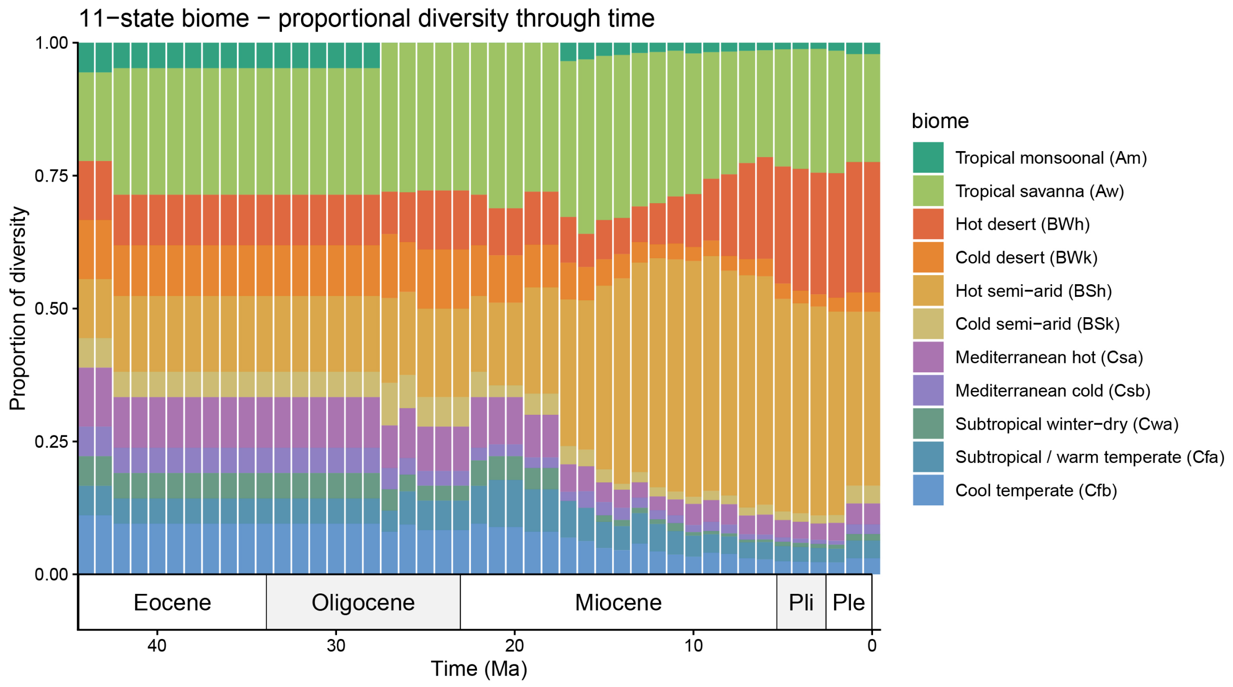

Figure S6. Paleobiome reconstructions used to inform BioGeoBEARS time-stratified DEC analysis. Stacked area chart showing proportional occupancy of all biomes through time.

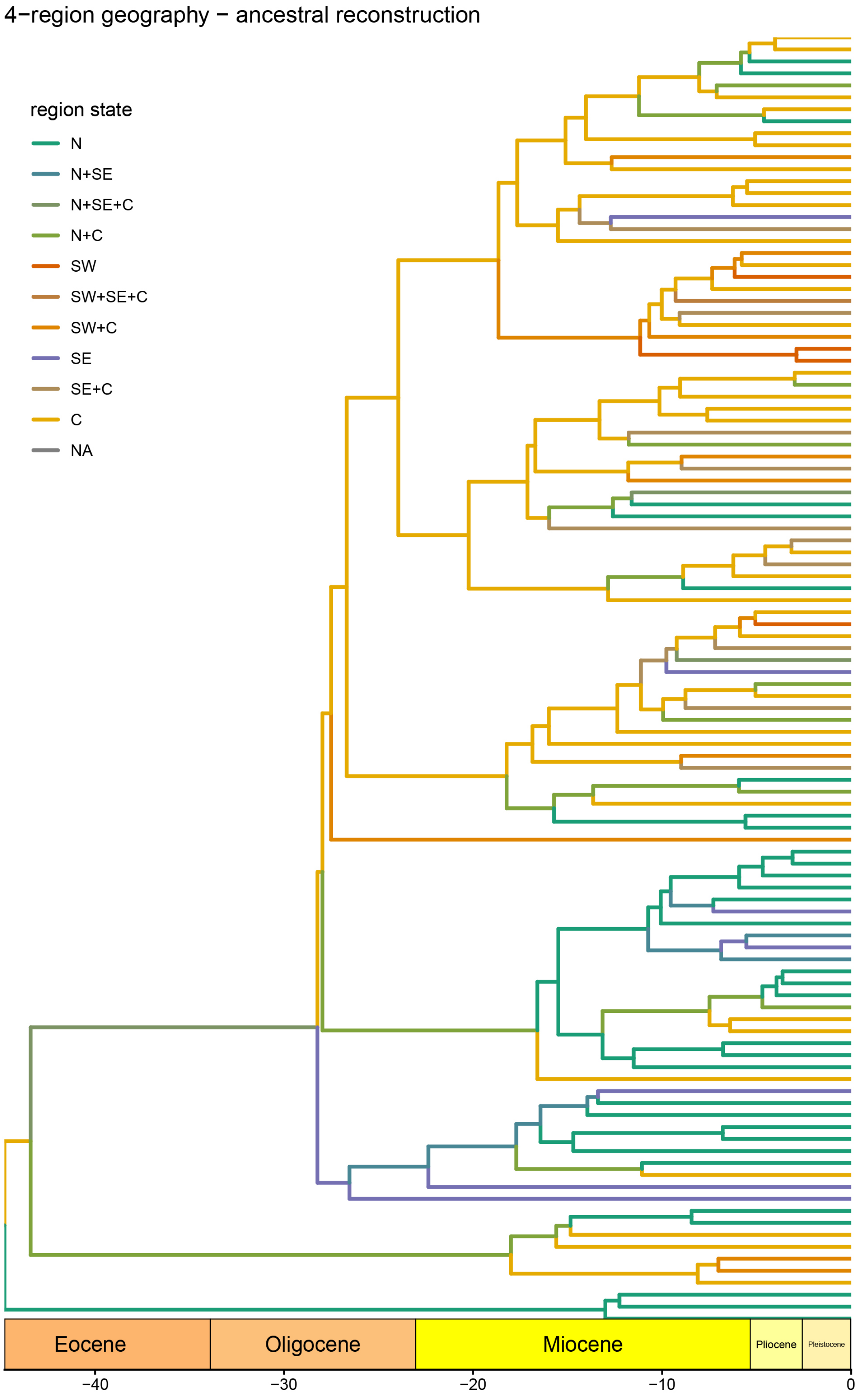

Figure S7. Ancestral range estimation based on four-region geography model inferred using BioGeoBEARS time-stratified DEC analysis.

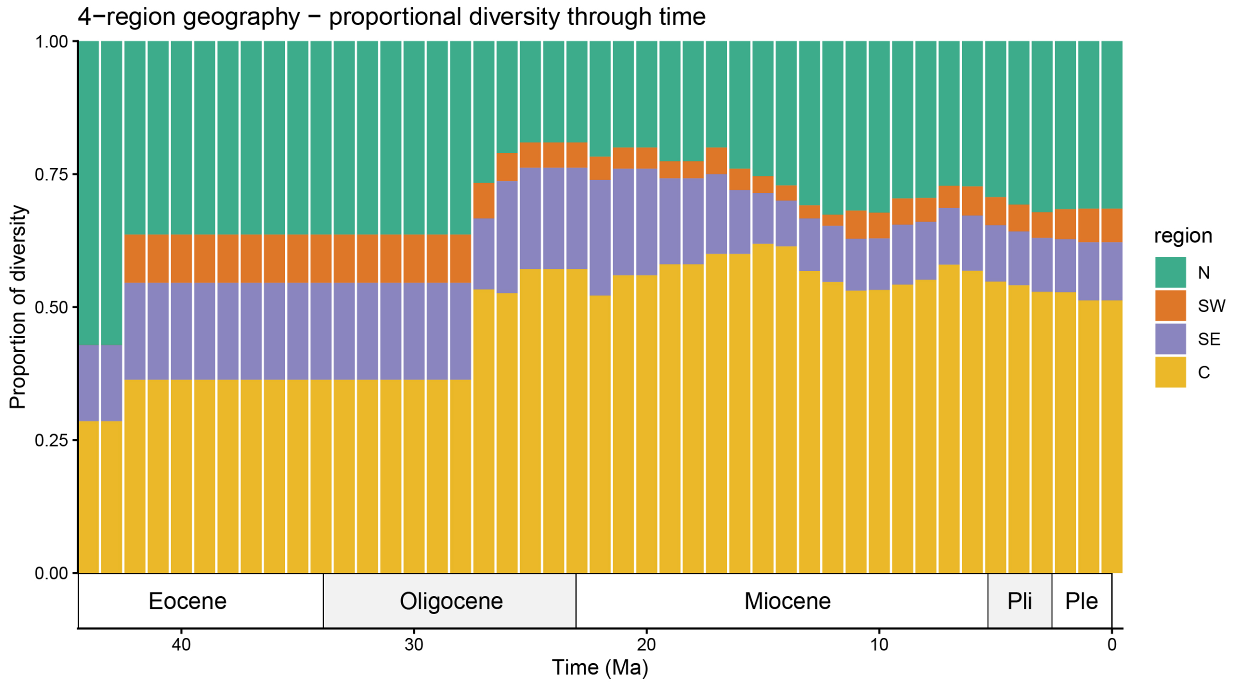

Figure S8. Stacked area chart showing proportional of diversity in the four regions through time

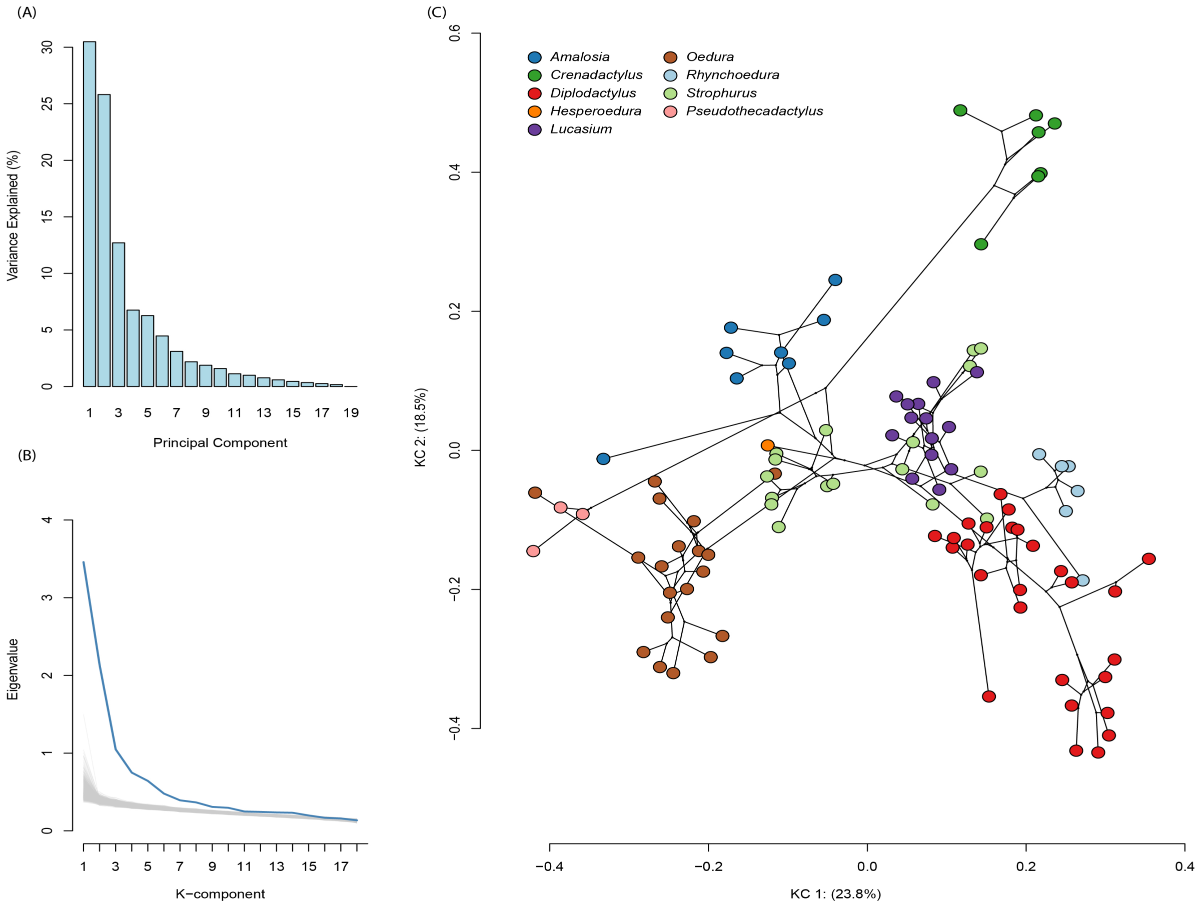

Figure S9. a) Scree plot showing variance explained by each principal component from the multivariate trait dataset (19 traits). b) Observed eigenvalues (blue line) obtained from multivariate phylogenetic signal matrix, K, along with 95% intervals of the permutation distributions in grey. Deviations from 95% intervals indicate the relative strength of phylogenetic signal for each K-component. c) First two K-components of diplodactylid log shape ratios. These represent the traits with maximal phylogenetic signal. Black lines represent phylogenetic relationships and estimated ancestral state according to Brownian Model of evolution. Tip colors represent different genera. The phylogenetic structure is much better represented by the first 2 K-components compared with the ordinary PCs (in main text)

### Supplementary Tables

Table S1. MCMCTree priors and calibrations used in genus-level and species-level divergence time analyses.

| ID | node | Split | Fossil name | Min. age | Max. age | MCMCTree prior 1 | Authorship use | Taxon description | Specimen ID |
|---|---|---|---|---|---|---|---|---|---|
| G1 | 23 | Lepidosauria + Archosauria | Protosaurus speneri | 254.7 | 269.3 | B(2.547,2.693) | Ezcurra et al. 2014 | von Meyer 1832 | RCSHC-Fossil Reptiles 308; NHMW 1943I4 |
| G2 | 24 | Sphenodon–Dibamus | Lepidosaurian Vellberg | 238.0 | 251.4 | B(2.475,2.550) | Jones et al. 2013 | Jones et al. 2013 | SMNS 91060; SMNS 91061 |
| G3 | 25 | Stem Gekkonomorpha | Eichstaettisaurus schroederi | 150.0 | 215.0 | B(1.500,2.380) | Simões et al. 2018 | Broili 1938 | MB.R.2035 |
| G4 | 40 | Stem Scincomorpha | Becklesius hoffstetteri | 152.1 | 200.0 | B(1.521,2.380) | Pyron 2016; Bolet et al. 2022 | Seiffert 1973 | |
| G5 | 42 | Serpents + Lizards | Eophis underwoodi | 166.1 | 200.0 | B(1.661,2.380) | Caldwell et al. 2015 | Caldwell et al. 2015 | |
| G6 | 43 | Varanids + Agamids | Dorsetisaurus sp. | 148.0 | 200.0 | B(1.480,2.380) | Jones et al. 2013 | Prothero and Estes 1980 | AMNH 27646, AMNH27647 |
| S1 | – | Crown Aotearoa New Zealand Diplodactylidae | Numerous see reference | | 16.0 | L(0.16,0.1,0.09) | Daza et al. 2014; Lee et al. 2009 | Lee et al. 2009 | |

Table S2. Fossil and secondary calibrations for species-level divergence time estimation.

| Node | Mean (normal) | SD (normal) | Skew-T prior |
|---|---|---|---|
| n23 | 262.0169 | 4.4269 | 'ST(2.6220361,0.0442471,-0.0525982,279.2273972)' |
| n24 | 252.0202 | 2.1864 | 'ST(2.5512903,0.0379655,-13.1002642,581.4362853)' |
| n25 | 208.1600 | 8.0550 | 'ST(2.0224247,0.0997224,1.106719,414.4988699)' |
| n26 | 127.0694 | 20.1162 | 'ST(1.1116272,0.2562405,1.2365927,589.5472223)' |
| n27 | 62.1637 | 10.6346 | 'ST(0.5061525,0.1529674,2.4953026,40.0442664)' |
| n28 | 47.9537 | 8.1765 | 'ST(0.3911833,0.1165724,2.4789744,32.6363785)' |
| n29 | 45.3786 | 7.8485 | 'ST(0.3697498,0.1107762,2.4219598,27.9847933)' |
| n30 | 42.8307 | 7.5471 | 'ST(0.3476849,0.1069059,2.3898422,32.1411399)' |
| n31 | 27.1392 | 4.8473 | 'ST(0.220732,0.067878,2.2426277,32.3935981)' |
| n32 | 26.0720 | 4.7239 | 'ST(0.2116876,0.0657741,2.2054093,30.4543997)' |
| n33 | 21.3859 | 4.2267 | 'ST(0.170422,0.0580719,2.1699677,24.840204)' |
| n34 | 15.5670 | 3.6897 | 'ST(0.1181504,0.0501096,2.1284818,21.8530403)' |
| n35 | 26.8455 | 4.8249 | 'ST(0.2182476,0.0673708,2.2168661,31.7767824)' |
| n36 | 25.5860 | 4.7424 | 'ST(0.2076836,0.0652368,2.0830526,29.4766448)' |
| n37 | 44.4529 | 7.9825 | 'ST(0.3616001,0.1104964,2.2313549,25.739724)' |
| n38 | 57.5713 | 10.3135 | 'ST(0.4665348,0.1460613,2.3112218,37.3535955)' |
| n39 | 202.3540 | 7.7640 | 'ST(1.9616157,0.0990332,1.2526228,342.596346)' |
| n40 | 193.0170 | 7.2734 | 'ST(1.8630918,0.0985531,1.6130442,247.4090008)' |
| n41 | 178.6001 | 6.0436 | 'ST(1.7271335,0.0796049,1.9318668,17.9343085)' |
| n42 | 168.0815 | 4.8483 | 'ST(1.643506,0.0480621,1.4268661,4.8658109)' |
| n43 | 156.7067 | 6.0836 | 'ST(1.5101672,0.0822224,1.6746968,78.9941116)' |

Table S3. Definition of morphological measurements used in comparative analyses.

| No | Measurement | Abbreviation | Method |
|---|---|---|---|
| 1 | Snout-vent length | SVL | From tip of the snout to the vent (cloaca) |
| 2 | Snout-axilla | SAL | From the tip of the snout to the midpoint of the crease between the forelimb and the body on the ventral surface |
| 3 | Interlimb length | ILL | Midpoint of the crease on the ventral surface where the fore-limb connects to the body, to the midpoint of the crease on the ventral surface where the hind-limb connects to the body |
| 4 | Body width | BW | From one lateral side of the body to the other at the midpoint of the ILL |
| 5 | Pelvic width | PW | From the midpoint of the crease on the ventral surface where the left hind limb connects to the body, to the midpoint of the crease on the ventral surface where the right hind limb connects to the body |
| 6 | Pelvic height | PH | From the top of the dorsal surface were the PW was measured, to the bottom of the ventral surface where the PW was measured |
| 7 | Head width | HW | Widest part of the head from one dorsal-lateral edge to the other edge |
| 8 | Head depth | HD | From the top of the tallest part of the head on the dorsal surface, to the bottom of the ventral surface under the jaw. |
| 9 | Head Length | HL | From the nose tip, to the anterior of the ear. |
| 10 | Snout length | SN | From the nasal opening to the anterior of the eye. |
| 11 | Eye diameter | ED | From one side of the eye to the other. |
| 12 | Lower front limb | LFL | Measured from the base of the lower fore limb to the juncture where the limb meets the front foot. |
| 13 | Upper front limb | UFL | Measured from the crease on the ventral surface where the fore limb connects to the body, to the end of the lower front limb. |
| 14 | Hand | FFOOT | From the base of the foot to the end of the longest toe (claw included). |
| 15 | Upper hind limb | UHL | Measured from the crease on the ventral surface where the hind limb connects to the body, to the end of the knee. |
| 16 | Lower hind limb | LHL | Measured from the top of the knee joint to the heel juncture where the limb meets the front foot. |
| 17 | Foot | HFOOT | From the base of the hind foot to the end of the longest toe (claw included) |
| 18 | Tail width | TW | Measured at the vent, from one dorsal-lateral edge to the other edge |
| 19 | Tail length | TL | Measured from the vent to the tip of the tail |

Table S4. BPP posterior clade probabilities in MAP species tree. Topology values show, for each combination of priors, the proportion of posterior trees that recovered the MAP topology versus all other topologies.

| Prior | $\theta$ Prior | $\tau$ Prior | MAP topology | Other topology |
|---|---|---|---|---|
| Set1 | 0.02 | 0.01 | 0.55500 | 0.4440 |
| Set2 | 0.02 | 0.03 | 0.77700 | 0.2220 |
| Set3 | 0.02 | 0.05 | 0.81045 | 0.1890 |
| Set4 | 0.10 | 0.01 | 0.77700 | 0.2220 |
| Set5 | 0.10 | 0.03 | 0.66660 | 0.3333 |
| Set6 | 0.10 | 0.05 | 0.77700 | 0.2220 |
| Set7 | 0.20 | 0.01 | 0.84189 | 0.1580 |
| Set8 | 0.20 | 0.03 | 0.66600 | 0.3330 |
| Set9 | 0.20 | 0.05 | 0.77760 | 0.2222 |

Table S5. Habitat type discrete character evolution model comparison. Models were fitted using equal-rates (ER), symmetrical (SYM), all-rates-different (ARD), and hidden-rate (HRM) Mk models, and an adjacency-constrained model.

| Model | log(L) | d.f. | AIC | weight |
|---|---|---|---|---|
| ER | -127.64 | 1 | 257.29 | 0.99 |
| SYM | -105.78 | 28 | 267.55 | 0.01 |
| ARD | -80.95 | 56 | 273.90 | 0.00 |
| ER + HRM | -174.92 | 3 | 355.83 | 0.00 |
| SYM + HRM | -174.08 | 57 | 462.15 | 0.00 |
| ARD + HRM | -132.16 | 114 | 492.33 | 0.00 |
| Adjacency model | -139.36 | 1 | 280.72 | 0.00 |

Table S6. Substrate orientation discrete character evolution model comparison. Models were fitted using equal-rates (ER), symmetrical (SYM), all-rates-different (ARD), and hidden-rate (HRM) Mk models, and an adjacency-constrained model.

| Model | log(L) | d.f. | AIC | weight |
|---|---|---|---|---|
| ER | -101.93 | 1 | 205.87 | 0.26 |
| SYM | -87.26 | 15 | 204.51 | 0.50 |
| ARD | -72.99 | 30 | 205.97 | 0.24 |
| ER + HRM | -133.32 | 3 | 272.63 | 0.00 |
| SYM + HRM | -133.32 | 31 | 328.63 | 0.00 |
| ARD + HRM | -123.59 | 62 | 371.19 | 0.00 |
| Adjacency model | -139.36 | 1 | 280.72 | 0.00 |

Table S7. Ecological state-dependent diversification model comparison (AIC). k is the total number of parameters. Turnover, extinction, and transition parameters are also shown. Character-independent models outperformed character-dependent models

| Model | logLik | k | turnover | extinction | transitions | AIC | ΔAIC | AICweight |
|---|---|---|---|---|---|---|---|---|
| CID3 | -578.17 | 11 | 3 | 1 | 7 | 1178.34 | 0.00 | 0.416 |
| CID4 | -577.34 | 12 | 4 | 1 | 7 | 1178.68 | 0.33 | 0.352 |
| CID5 | -577.19 | 13 | 5 | 1 | 7 | 1180.38 | 2.04 | 0.150 |
| CID6 | -577.20 | 14 | 6 | 1 | 7 | 1182.41 | 4.06 | 0.054 |
| CID2 | -581.84 | 10 | 2 | 1 | 7 | 1183.69 | 5.35 | 0.029 |
| MuHiSSE | -580.46 | 20 | 6 | 1 | 13 | 1200.92 | 22.58 | 0.000 |
| Null MuSSE | -596.20 | 8 | 1 | 1 | 6 | 1208.40 | 30.06 | 0.000 |
| Full MuSSE | -595.34 | 10 | 3 | 1 | 6 | 1210.68 | 32.33 | 0.000 |

Table S8. Speciation ($\lambda$), extinction ($\mu$), and net diversification (r) rates under the best-supported MuHiSSE model. Values show mean (95% CI) for each ecological state.

| Rate | Open | Arboreal | Saxicolous |
|---|---|---|---|
| $\lambda$ (speciation rate) | 0.115 (0.104–0.126) | 0.138 (0.125–0.151) | 0.14 (0.126–0.152) |
| $\mu$ (extinction rate) | 2.37e-10 (2.15e-10–2.6e-10) | 2.85e-10 (2.56e-10–3.12e-10) | 2.88e-10 (2.6e-10–3.14e-10) |
| r (diversification rate) | 0.115 (0.104–0.126) | 0.138 (0.125–0.151) | 0.14 (0.127–0.152) |

Table S9. Univariate phylogenetic GLS results for body size (geometric mean) among ecological states. LCL/UCL are 95% confidence limits. Δ vs. ground shows the difference relative to ground-dwelling lineages. P-values < 0.05 are shown in blue.

| Ecological state | Mean | SE | LCL | UCL | Δ vs. ground | SE (Δ) | P-value |
|---|---|---|---|---|---|---|---|
| Ground | 1.86 | 0.12 | 1.63 | 2.09 | | | NA |
| Rock | 2.24 | 0.11 | 2.02 | 2.47 | 0.38 | 0.10 | 0.003 |
| Spinifex | 1.79 | 0.14 | 1.52 | 2.06 | -0.07 | 0.12 | 0.95 |
| Spinifex Ground | 1.54 | 0.13 | 1.28 | 1.80 | -0.32 | 0.10 | 0.007 |
| Tree | 2.19 | 0.11 | 1.97 | 2.41 | 0.33 | 0.10 | 0.008 |
| Tree Ground | 1.92 | 0.17 | 1.57 | 2.27 | 0.06 | 0.17 | 0.99 |
| Tree Rock | 2.29 | 0.11 | 2.07 | 2.51 | 0.43 | 0.10 | 0 |

Table S10. Univariate phylogenetic GLS results for tail length among ecological states. LCL/UCL are 95% confidence limits. Δ vs. ground shows the difference relative to ground-dwelling lineages. P-values < 0.05 are shown in blue.

| Ecological state | Mean | SE | LCL | UCL | Δ vs. ground | SE (Δ) | P-value |
|---|---|---|---|---|---|---|---|
| Ground | 1.57 | 0.14 | 1.29 | 1.84 | | | NA |
| Rock | 1.69 | 0.13 | 1.42 | 1.95 | 0.12 | 0.12 | 0.78 |
| Spinifex | 1.66 | 0.16 | 1.34 | 1.98 | 0.10 | 0.15 | 0.92 |
| Spinifex Ground | 1.67 | 0.15 | 1.37 | 1.97 | 0.10 | 0.11 | 0.79 |
| Tree | 1.79 | 0.13 | 1.54 | 2.05 | 0.22 | 0.12 | 0.24 |
| Tree Ground | 1.40 | 0.20 | 1.00 | 1.81 | -0.16 | 0.19 | 0.85 |
| Tree Rock | 1.76 | 0.13 | 1.50 | 2.01 | 0.19 | 0.12 | 0.42 |

### Supplementary Methods

#### Taxonomic sampling and sequencing

We sequenced 190 individuals representing 108 of the 111 recognized species of Australian diplodactylids. We sourced tissues that were frozen or stored in ethanol from the South Australian Museum, Queensland Museum, Museum and Art Gallery of the Northern Territory, Western Australian Museum, and from private collections. Our sampling did not include Diplodactylus kenneallyi, known only from the single type specimen, and Strophurus congoo and S. trux, for which no tissues were available. Sample preparation and sequencing methods followed in Tiatragul et al. (2023a).

In brief, we extracted DNA using the Macherey-Nagel nucleospin extraction kit at the Australian National University (ANU). We fragmented and selected the fragments over 600 bp. We prepared libraries using NextFlex Rapid NDA-Seq Kit 2.0 (PerkinElmer) and barcoded each fragment with Unique Dual Index Barcode and enriched our libraries using six cycles of PCR. We targeted nuclear exons with the custom designed Squamate Conserved Loci (SqCL) probes representing 388 anchored hybrid enrichment (AHE) loci, 5,023 ultraconserved elements (UCEs) and 42 nuclear legacy exons (NLE) (Singhal et al. 2017). The hybridized libraries were sequenced on an Illumina HiSeq 2500 platform at the ANU.

We assembled and aligned the sequences using the Nextflow pipeline pipesnake v.1.2 (Brennan et al. 2024b). We used the `--end-prg` argument to stop pipesnake at the pseudo-reference genome (PRG) step. We specified the PRGs we wanted to include and ran the pipesnake pipeline with the `--from-prg` argument. At this stage, the program creates a locus-specific raw alignment that includes targets from all the samples before performing multi sequence alignment (MSA) with MAFFT v.7 (Katoh and Standley 2013).

To expand our taxonomic sampling of other diplodactylids, we included species from NC ($n = 31$) and ANZ ($n = 18$) by incorporating published sequence data (Skipwith et al. 2019; Title et al. 2024). To ensure these sequences are orthologous, we matched the dataset to common Squamate Conserved Loci (SqCL) targets (Singhal et al. 2017) and renamed each locus. The post-processed raw alignment dataset includes 5,453 loci comprising 383 AHE loci, 4,995 UCEs, and 37 NLEs. Next, we trimmed the raw MSA outputs from pipesnake with ClipKit v.2.3.0 using the 'smart-gap -m gappy 0.8' (Steenwyk et al. 2020). Additionally, we used the correction_multi_aggressive.jl program in TAPER v.1.0.0 (Zhang et al. 2021)—written in Julia (Bezanson et al. 2012)—to identify and mask outliers in the trimmed alignment sequences given the overall level of divergence of the sequence to others (Zhang et al. 2021). From the full-sample (n = 276 taxa) trimmed and TAPER-corrected alignments, we subsampled 171 diplodactylid lineages plus Sphenodon punctatus, Phyllurus ossa, and Aprasia aurita as outgroups for divergence dating, and nine taxa for the genus-level species tree analysis (Extended Data 01). Each of these datasets were filtered, trimmed and TAPER-corrected separately.

#### Phylogenetic inference

We applied two phylogenetic inference approaches to estimate the species tree from the trimmed and aligned SqCL dataset. These methods are discussed in detail below, but briefly: the first approach was a weighted summary coalescent method to estimate the species tree from the gene trees, and the second was a full Bayesian multispecies coalescent (MSC) model sampling a single representative from each genus to estimate higher-level relationships.

##### Summary coalescent method and discordance analyses

We estimated maximum likelihood gene trees for each SqCL locus using IQ-TREE v2.3.6 (Minh et al. 2020), using ModelFinder to assign the best-fitting molecular evolution model and subsequently perform 1000 ultrafast bootstrap approximation (UFBoot) replicates to estimate branch support. The output unrooted gene trees were used as input for weighted ASTRAL (wASTRAL) v1.22.3.7 (Zhang and Mirarab 2022) to infer a species tree with the `-u 2` flag used to calculate alternative "quartet scores" and branch local posterior probability. wASTRAL accounts for phylogenetic uncertainty by integrating signals from branch length and branch support in gene trees.

To quantify phylogenetic concordance and discordance, we used the concordance vector ($\Psi$) framework, which incorporates gene, site, and quartet concordance factors (Lanfear and Hahn 2024). We calculated the gene concordance factor (gCF) and site concordance factor (sCF)—the proportion of "decisive" gene trees/sites for which a given branch in the reference species tree is true—using IQ-TREE (Minh et al. 2020) the topology fixed to the wASTRAL species tree. Site concordance factors were calculated using the likelihood calculation option (`-scfl`) (Mo et al. 2023). The quartet frequencies from wASTRAL are treated as quartet concordance factors (qCFs) as they indicate the proportion of relevant quartets associated with the topology assigned for the particular branch of interest (Lanfear and Hahn 2024).

Inferring the Australian diplodactylid species tree using an MSC model

The placement of Hesperoedura within the core Australian clade is uncertain (Oliver et al. 2012; Skipwith et al. 2019); as a potentially early-branching lineage confined to relictual mesic forests of southwestern Australia, its position carries direct implications for ancestral biome reconstruction. We therefore estimated an alternative species tree for the core Australian diplodactylids using a full Bayesian MSC model in BPP v.4.7.0 (Yang 2015; Flouri et al. 2018) on the genus-level dataset (4,458 loci; 9 taxa) including: Amalosia rhombifer, Diplodactylus vittatus, Hesperoedura reticulata, Lucasium alboguttatum, Oedura elegans, Rhynchoedura angusta, Strophurus intermedius, and Crenadactylus horni as the outgroup(Oliver and Sanders 2009). The MSC model involves two parameters: node height ($\tau$) and the effective population size ($\theta = 4N\mu$). We first ran an 'A00' analysis (speciesdelimitation = 0; speciestree = 0) with a fixed species tree based on the topology from wASTRAL to estimate these parameters, using diffused inverse-gamma (InvG) priors to both parameters, with a shape of 3 for each and with one of three different mean values representing the maximum, mean, and minimum approximated from the posterior in preliminary runs.

We then ran nine sets of 'A01' analyses to estimate species tree with different $\theta$ and $\tau$ priors: $\theta \sim$ InvG(3, 0.02), InvG(3, 0.10), and InvG(3, 0.20). For each $\theta$ prior, we assigned a corresponding $\tau$ prior of InvG(3, 0.01), InvG(3, 0.03), and InvG(3, 0.05). Population size parameters ($\theta$) are integrated out analytically to improve mixing of the MCMC, while $\tau$ at non-root nodes were generated from a uniform-Dirichlet distribution (Yang 2015). For each set we ran the MCMC for 400,000 iterations after burn-in of 20,000 iterations, with samples taken every second iterations (nsample = 200,000). We repeated each A01 analysis set nine times using different starting species trees and assessed convergence by comparing posterior topology distributions across runs. Of the 81 total runs, we discarded 20 that did not give a dominant topology. We combined the remaining 61 runs to produce the species tree with the highest posterior probability (i.e., the maximum a posteriori tree — MAP tree). We generated a maximum clade credibility (MCC) tree from the combined samples using TreeAnnotator v2.7.5 (Bouckaert et al. 2014). We then performed a final set of A00 analysis with the species tree fixed at the MAP tree to estimate the $\theta$ and $\tau$ parameters (Rannala and Yang 2017; Flouri et al. 2018). The prior used in the A00 analyses were $\theta \sim$ InvG(3, 0.1) for all populations and $\tau \sim$ InvG(3, 0.03). We ran the MCMC for 1,000,000 iterations after burn-in of 100,000 taking samples every 10 iterations (nsample = 100,000). We repeated this A00 analysis was repeated eight times, checked for consistency of posterior summaries and combined all eight.

Divergence time estimation

We estimated divergence times using MCMCTree in the PAML package (Yang 2007). Divergence dating was based on four partitioned AHE and one unpartitioned UCE datasets, with species-level topology fixed to match the wASTRAL phylogeny. To incorporate fossil calibrations, we added our AHE alignments to an alignment with nine reference lineages (one bird and eight reptiles) from Brennan et al. (2024a). Raw genetic distances from this reference alignment were used as proxies for evolutionary rate. To minimize issues from extreme rate heterogeneity, we discarded the fastest and slowest 5% of loci, retaining 388 AHE loci and partitioning them into four rate categories (i.e., "GeneClumps"). The four rate categories correspond to the four quartiles of rate variation. From each AHE locus, we used AMAS (Borowiec 2016) to retain codon positions 1 and 2 and discarded the third positions. For the unpartitioned UCE dataset, we selected the 100 best-ranking UCE loci based on a "gene shopping" approach following (Smith et al. 2018) using a custom R script from (Borowiec et al. 2025). We identified a subset of loci that had desirable properties of clock-likeness (low root-to-tips variance), high information content (high average gene tree branch lengths), and low topological discordance (low Robinson-Foulds (RF) distance) with the reference species tree by computing each metric and ranking each loci. We used the sum of weighted ranks to obtain the most desirable loci. Weighted ranks were 0.5 for clock-likeness, 0.3 for average branch lengths, and 0.2 for RF distances. Loci within each partition were concatenated, and SEGUL v.0.22.1 (Handika and Esselstyn 2024) was used to extract the relevant taxa and convert alignments into PHYLIP format for MCMCTree.

For each partition, we estimated approximate likelihoods and branch lengths using baseml with `usedata = 3` (dos Reis and Yang 2011). The resulting .BV files were concatenated, and MCMCTree was run using the gradient and Hessian matrix (usedata = 2). Each run consisted of a burn-in of 10,000 generations followed by sampling every 100 generations until 10,000 samples were collected (1,010,000 total generations). Convergence and stationarity were assessed by ensuring ESS $> 200$ for all parameters. Four runs were combined using LogCombiner (Bouckaert et al. 2019), and divergence times were summarized on the species tree (print = -1 in.ctl file). We conducted two analyses, a preliminary, genus-level analysis including one representative per genus, calibrated with six squamate fossils (Table S1). Second, a full species-level analysis using five fossil calibrations and 13 secondary calibrations derived from the posterior of the genus-level run (Table S2).

#### Statistical analyses

##### Paleobiome reconstruction and historical biogeography

To estimate the biogeographic history of the clade across major biomes on the Australian continent we fitted a dispersal-extinction-cladogenesis (DEC) model with the R package BioGeoBEARS v.1.1.3 (Matzke 2013). We used the Köppen-Geiger climate classification as a biome scheme for this analysis as this classification scheme is based on monthly variation in temperature and precipitation values and is transferable through deep time, which is appropriate for ectotherms. First, we estimated the occurrence of each species of diplodactylid in each biome in Australia by overlaying occurrence records with a present-day model of Köppen-Geiger biomes (Beck et al. 2018). Biomes are classified using letters with the first letter defining the broad biome (A = tropical, B = arid, C = temperate, D = cold, and E = polar), with additional letters determining the seasonality of temperature and precipitation (for example "Af" is tropical rainforest and "Aw" is tropical savanna). To reduce the absolute number of states in the model, we only classed species as belonging to a biome if they had at least 10% of their records in a biome, allowing a maximum number of ranges occupied to be four. Biomes have shifted their distributions dramatically across Australia during the period of diplodactylid diversification, with some like the dunefields of the arid zone appearing very late in the groups history (Pepper and Keogh 2021). To incorporate realistic constraints on lineage dispersal imposed by this paleoenvironmental history, we fitted a time-stratified DEC model that constrained dispersal probabilities and the available state space according to biome availability through time.

We estimated Köppen-Geiger biomes from a general circulation model (GCM) climatology of Li et al. (2022) at 10 Ma intervals from the present to 50 Ma following the method of Peel et al. (2007), after first down-scaling the GCM data from coarse spatial resolution (1 degree) to high resolution (0.1 degree) using the CHELSA down-scaling algorithm (Karger et al. 2017) and 0.1 degree topographical template (Scotese and Wright 2018). This ensured we were able to capture the fine scale distribution of biomes due to topography that are lost at coarser resolution. In addition to constraining the state space at 10 Ma time intervals, we also modified dispersal probabilities between regions based on geographic distances estimated as the minimum pairwise distances between biomes at each time step. Finally, we also constrained dispersal by the environmental distances between biomes by first extracting 19 Bioclimatic variables across diplodactylid occurrence records taken from CHELSA (Karger et al. 2017). We performed a PCA to reduce the dimensionality and collinearity of these variables to a set of orthogonal axes and obtained the pairwise distance between points from each biome in this climate space. The fitted DEC model included additional free parameters n which weighted the effect of environmental distances and x which weighted the effect of geographic distances.

Finally, to investigate whether the biome classification led to qualitative differences in estimating the clades ancestral areas, we fitted a geography-only DEC model. In that model we categorized species as occupying the major geographic regions of Australia: Southwest, Southeast, Northern, and Central. We qualitatively compared the results between climatic-ecoregion and geography-only models.

##### Morphological association with ecological state and phylogenetic signal

To characterize the major axes of morphological variation, we conducted a principal component analysis (PCA) on the full 19-trait dataset using the `PCA` function in the R package FactoMineR v.2.11. We retained 12 functionally informative traits for subsequent comparative analyses, selected based on their contribution to the major PCA axes and functional relevance across six morphological modules: tail (tail length, tail width), body (body width, pelvic height), forelimb (upper and lower arm length), hindlimb (upper and lower leg length), head (head depth, posterior skull length), and neck length, plus geometric mean of size. PCA scores were used to visualize morphological differentiation among ecological states and to summarize multivariate patterns of shape variation. This two-step approach allowed exploratory identification of major axes of morphological variation followed by hypothesis testing on a reduced set of functionally informative traits.

To test whether ecological state explains variation in morphology among Australian diplodactylids, we fitted multivariate phylogenetic generalized least squares models using `mvgls` in the R package mvMORPH v.1.2.1 (Clavel et al. 2015). The 12-trait subset were analyzed jointly as a multivariate response, with ecological state specified as a fixed effect and phylogenetic relatedness incorporated through the residual covariance structure using Pagel's .

Significance of ecological predictors was evaluated using permutation-based multivariate analysis of variance (MANOVA) with Pillai's trace and 1000 permutations. We additionally tested morphological associations with substrate orientation in separate models to assess alternative ecological sources of morphological variation. Where overall ecological effects were significant, pairwise post-hoc comparisons among ecological states were conducted using the `pairwise.glh` function based on 1000 permutations on Pillai's test statistic and p-value adjusted using BH correction (Benjamini and Hochberg 1995). To test directional predictions, we additionally fitted univariate phylogenetic generalized least squares models for size and tail traits with ecological state as a fixed effect.

We quantified phylogenetic signal in morphological variation to assess the degree to which trait similarity among species reflects shared ancestry versus ecological differentiation. Phylogenetic signal was estimated using a multivariate extension of Blomberg's K, which assesses whether closely related species resemble each other more than expected under a Brownian motion model of evolution (Blomberg et al. 2003; Adams 2014). To evaluate whether phylogenetic signal was uniformly distributed across trait dimensions or concentrated along specific axes of variation, we additionally calculated the arithmetic mean of eigenvalues of K ($K_A$) and geometric mean of the eigenvalues of K ($K_G$) (Mitteroecker et al. 2025).

##### Posthoc: Morphological association with hidden diversification rate classes

To test whether morphological variation corresponds to the hidden rate classes identified by the best-supported diversification model, we extracted tip-level marginal posterior probabilities for each hidden rate class from the CID3 marginal reconstruction. We obtained posterior probabilities for each species across the three hidden classes by summing state probabilities across observed ecological states within each hidden class. We assigned species to either low- or high-diversification-rate groups based on their dominant posterior probability, which revealed an effectively bimodal structure in class-level mean speciation rates. The two lower-rate classes ( 0.048 My$^{1}$) were pooled into a single slow group for primary analyses, while the elevated class ( 0.073 My$^{1}$) was designated as the fast group (n = 75 slow, n = 84 fast). As a complementary approach, we also used the continuous posterior probability of belonging to the high-rate group as a predictor, which retains the uncertainty of the marginal reconstruction without requiring hard assignment.

We tested whether body size, tail length, and tail width — the traits with strongest ecological and phylogenetic signal in our morphological analyses — differed between high- and low-rate groups using univariate phylogenetic generalized least squares models (Pagel's , same specification as the ecological morphology models described above). We additionally tested the full 12-trait matrix using multivariate phylogenetic GLS (mvgls, mvMORPH v.1.2.1) with permutation-based MANOVA (Pillai's trace, 1000 permutations). We restricted this analyses to species with complete morphological measurements (n = 102).

### Supplementary Results

#### Posthoc: Morphological association with hidden diversification rate classes

Tip-level posterior probabilities from the CID3 marginal reconstruction showed a bimodal rate structure with 75 species assigned to the low-rate group (mean = 0.048 My$^{1}$) and 84 to the high-rate group (mean = 0.073 My$^{1}$), and only one assigned to a third class at the boundary. None of the univariate phylogenetic GLS models were significant: body size (F= $< 0.001$, df = 100, $p > 0.99$), tail length (F = = 1.37, df = 100, p = 0.24), and tail width (F = 0.38, df = 100, p = 0.54) did not differ between rate groups. Results were consistent when the continuous posterior probability of high-rate group membership was used as a predictor (all $p > 0.10$). The multivariate phylogenetic MANOVA on the full 12-trait matrix was similarly non-significant (Pillai's trace = 0.099, p = 0.498). These results indicate that the hidden diversification rate variation identified by CID3 is not explained by any of the morphological traits measured here.

### Supplementary Discussion

#### Ancestral biome

Biogeographic reconstruction provides a complementary perspective on how environmental change structured the spatial and ecological assembly of the clade. BioGeoBEARS analyses identify the seasonal tropics of northern Australia as the most likely ancestral biome for Australian diplodactylids, followed by expansion into semi-arid and ultimately fully arid biomes. This pattern is consistent with directional expansion into progressively drier environments in which the radiation originated in relatively mesic environments and subsequently tracked the geographic expansion of open and increasingly dry habitats through time (Byrne et al. 2011). Notably, modern Australian diplodactylids are largely absent from rainforest environments except for a small number of species in Pseudothecadactylus and Amalosia, whereas rainforest systems in NC and ANZ retain predominantly arboreal diplodactylid assemblages. In Australia, rainforest gecko communities are instead dominated by the closely related Carphodactylidae (Brennan and Oliver 2017), suggesting long-term ecological partitioning among Pygopodoidea lineages across biome types.

The inferred history of biome occupancy is highly dynamic, with an average of 202 transitions across the phylogeny, with over a third of these into desert environments. A major shift into arid environments was inferred for the Rhynchoedura–Diplodactylus–Lucasium clade near the Miocene onset, coinciding with continental drying — a pattern paralleled in agamid lizards, Pygopodoidea, and typhlopid snakes (Brennan and Oliver 2017; Tiatragul et al. 2023b; Brennan et al. 2025), many of which exhibit parallel transitions from mesic to increasingly arid biomes during the Miocene.

Despite clear directional expansion into arid environments, ancestral biome reconstruction reveals substantial uncertainty at deeper nodes, reflecting a complex history of repeated biome transitions rather than long-term biome conservatism. Many extant arid-zone lineages trace their origins to ancestors inferred to have occupied mesic or semi-arid environments prior to the full establishment of the modern arid zone. The crown age of the large predominantly arid clade (Strophurus, Diplodactylus, Lucasium, Rhynchoedura) dates to the Oligocene, preceding the widespread development of Australia's interior deserts. While this period, post Eocene-Oligocene Boundary, marks the onset of major drying in Australia (Byrne et al. 2008), much of the truly arid zone is thought to only emerge towards the late Miocene, with the development of dunefields as recently as the late Pleistocene ~1 Mya (Pepper and Keogh 2021). Consequently, many arid-zone species in the phylogeny are estimated to have diverged before the arid zone they occupy existed—a pattern that has been found in multiple arid lineages in Australian taxa (Skeels et al. 2025), and represents a major puzzle in Australian biogeography. Our paleoenvironmental reconstructions point towards a possible explanation for this: semi-arid climates are reconstructed to have existed across large parts of northern Australia throughout the Cenozoic while much of the rest of the continent, including inland Australia, was mesic. It is possible that early diverging lineages of the Diplodactylidae occupied this semi-arid biome leading to preadaptation to the increasing aridification that came later. Very little fossil or lithological proxy data is available for northern Australia and therefore it is poorly understood what the degree of early aridification was in this part of the continent. However, multiple climate models independently point towards a much drier climate, and this warrants further attention to understand the assembly of Australia's arid zone biota.The geography-only model showed that the most likely location for ancestral diplodactylids was in northern and central parts of Australia, pre-desertification of the continent. This could suggest that microhabitat affinities not captured by broad scale climatic classifications may be important in defining species distributions over deep time scales.

Taken together, the temporal mismatch between lineage origin and biome formation might explain the large number of inferred biome transitions during the Miocene. As aridity intensified and biome boundaries shifted across the continent, lineages moved through a series of transitions from tropical, to semi-arid, to arid and repeatedly reorganized their geographic ranges in response to changing environmental conditions. Rather than reflecting instability in biome association, this pattern may suggest progressive tracking of shifting climatic and biome boundaries over time. In particular, ground-dwelling clades appear to have expanded across progressively drier landscapes as open environments spread, suggesting that ecological traits associated with terrestrial life in open habitats may have facilitated persistence in increasingly arid conditions.

Large-scale landscape structure may also have influenced how diplodactylids responded to continental drying. Much of central Australia is characterized by low topographic relief and extensive flat landscapes, in contrast to the more structurally complex forested or montane systems (Hesse 2010; Pepper and Keogh 2021). The relatively flat and low-relief landscape may have facilitated the widespread expansion of ground-dwelling lineages by reducing physical constraints associated with vertical substrate use and enabling occupation of broad, continuous terrestrial surfaces. In relatively flat landscapes, environmental heterogeneity is often generated by broad climatic gradients and variation in substrate or soil properties rather than by strong topographic barriers potentially promoting climatic rather than topographic drivers of population fragmentation (Stein et al. 2014). The diversification of ground-dwelling clades (Diplodactylus, Rhynchoedura, and Lucasium) may reflect not only the spread of open habitats but also the spatial continuity of low-relief environments that allowed terrestrial ecological strategies to be deployed across large portions of the continent.

Because biogeographic models are estimated from extant lineages, an additional factor that may contribute to the high number of inferred biome transitions is extinction associated with climatic instability, particularly in expanding arid environments. Fossil and phylogenetic evidence indicates that Australia's arid zone has experienced repeated contraction and expansion throughout the Neogene, likely resulting in substantial lineage turnover (Byrne et al. 2008; Brennan and Oliver 2017; Pepper and Keogh 2021). If intermediate or transitional populations were lost during these cycles, ancestral reconstructions based on extant taxa would tend to infer more direct biome shifts than actually occurred. Under this scenario, the large number of inferred transitions may partly reflect the loss of lineages that formerly occupied intermediate environmental states rather than repeated discrete biome shifts. Instead of contradicting the pattern of dynamic biome occupancy, such extinction-driven pruning would reinforce the interpretation that lineage distributions were repeatedly reorganized in response to environmental change.

Together, these interacting processes indicate that continental aridification did not simply produce a static new biome that lineages subsequently invaded. Instead, environmental change generated a moving mosaic of habitat types that repeatedly reorganized species distributions over millions of years. Directional expansion into drier environments, extinction-mediated lineage turnover, and the spatial continuity of low-relief landscapes together shaped the geographic and ecological assembly of the Australian diplodactylid radiation.

#### Divergence time estimates

Our divergence time estimates are younger than those reported in Skipwith et al. (2019), reflecting two main differences. First, Skipwith et al. (2019) calibrated the pygopodoid crown with a prior bounded between 50–70 Ma derived from multilocus mtDNA and nuclear exon studies (Oliver and Sanders 2009; Skipwith et al. 2016). Despite this prior, they reported a posterior estimate of 76 Ma (95% HPD: 65–86 Ma). By excluding this prior, we allow the pygopodoid crown (60 Ma; 95% HPD: 51–70 Ma) to be better informed by the data and our primary fossil calibrations. Second, Skipwith et al. (2019) used 250 UCEs selected for taxon completeness, whereas ours used 388 partitioned AHE loci and 100 UCEs chosen for clock-likeness and low topological discordance via gene shopping; these dataset differences affect rate estimates and consequently node ages. Topological differences between studies are likewise concentrated in deeper backbone relationships, particularly the placement of Hesperoedura reticulata, and likely reflect differences in genetic dataset composition. Low gCF and sCF values at the deepest backbone nodes confirm that uncertainty at these nodes persists regardless of dataset.

#### Tail morphology

A limitation in our analysis is that we only captured tail morphology through relative linear measurements. Diplodactylid tails vary dramatically in overall form, ranging from slender and tapering to markedly bulbous or structurally specialized configurations, reflecting substantial three-dimensional variation in shape (Green et al. 2024). Such complexity is unlikely to be fully represented by simple measures of relative length or width. Future studies incorporating 2- and 3-dimensional landmark-based geometric morphometric approaches like Green et al. (2024) across the full diplodactylid radiation would provide a more complete characterization of tail shape and may reveal functional and ecological associations that are not detectable using linear trait data alone.

Bouckaert R., Vaughan T.G., Barido-Sottani J., Duchêne S., Fourment M., Gavryushkina A., Heled J., Jones G., Kühnert D., Maio N.D., Matschiner M., Mendes F.K., Müller N.F., Ogilvie H.A., Plessis L. du, Popinga A., Rambaut A., Rasmussen D., Siveroni I., Suchard M.A., Wu C.-H., Xie

D., Zhang C., Stadler T., Drummond A.J. 2019. BEAST 2.5: An advanced software platform for Bayesian evolutionary analysis. PLOS Computational Biology. 15:e1006650.

Brennan I.G., Chapple D.G., Keogh J.S., Donnellan S. 2024a. Evolutionary bursts drive morphological novelty in the world's largest skinks. Current Biology. 34:3905–3916.e5.

Brennan I.G., Cooper N., Sumner J., Tiatragul S., Tedeschi L.G., Broady E.S., Melville J., Keogh J.S. 2025. Generalists link peaks in the shifting adaptive landscape of Australia's dragon lizards. :2025.08.27.672751.

Brennan I.G., Oliver P.M. 2017. Mass turnover and recovery dynamics of a diverse Australian continental radiation. Evolution. 71:1352–1365.

Brennan I.G., Singhal S., Al Bkhetan Z. 2024b. Pipesnake: Generalized software for the assembly and analysis of phylogenomic datasets from conserved genomic loci. Bioinformatics. 40:btae195.

Byrne M., Steane D.A., Joseph L., Yeates D.K., Jordan G.J., Crayn D., Aplin K., Cantrill D.J., Cook L.G., Crisp M.D., Keogh J.S., Melville J., Moritz C., Porch N., Sniderman J.M.K., Sunnucks P., Weston P.H. 2011. Decline of a biome: Evolution, contraction, fragmentation, extinction and invasion of the Australian mesic zone biota. Journal of Biogeography. 38:1635–1656.

Byrne M., Yeates D.K., Joseph L., Kearney M., Bowler J., Williams M.A.J., Cooper S., Donnellan S.C., Keogh J.S., Leys R., Melville J., Murphy D.J., Porch N., Wyrwoll K.-H. 2008. Birth of a biome: Insights into the assembly and maintenance of the Australian arid zone biota. Molecular Ecology. 17:4398–4417.

Clavel J., Escarguel G., Merceron G. 2015. mvMORPH: An R package for fitting multivariate evolutionary models to morphometric data. Methods in Ecology and Evolution. 6:1311–1319.

dos Reis M., Yang Z. 2011. Approximate Likelihood Calculation on a Phylogeny for Bayesian Estimation of Divergence Times. Molecular Biology and Evolution. 28:2161–2172.

Flouri T., Jiao X., Rannala B., Yang Z. 2018. Species Tree Inference with BPP Using Genomic Sequences and the Multispecies Coalescent. Molecular Biology and Evolution. 35:2585–2593.

Green A.L., Oliver P.M., Gray J.A., Sherratt E. 2024. Adaptive tails? Parallel evolution of expanded tails in monsoonal tropics lineages of an Australian gecko radiation (Oedura). Zoological Journal of the Linnean Society. 202:zlad186.

Handika H., Esselstyn J.A. 2024. SEGUL: Ultrafast, memory-efficient and mobile-friendly software for manipulating and summarizing phylogenomic datasets. Molecular Ecology Resources. 24:e13964.

Hesse P.P. 2010. The Australian desert dunefields: Formation and evolution in an old, flat, dry continent. Geological Society, London, Special Publications. 346:141–164.

Karger D.N., Conrad O., Böhner J., Kawohl T., Kreft H., Soria-Auza R.W., Zimmermann N.E., Linder H.P., Kessler M. 2017. Climatologies at high resolution for the earth's land surface areas. Sci Data. 4:170122.

Katoh K., Standley D.M. 2013. MAFFT Multiple Sequence Alignment Software Version 7: Improvements in Performance and Usability. Molecular Biology and Evolution. 30:772–780.

Lanfear R., Hahn M.W. 2024. The Meaning and Measure of Concordance Factors in Phylogenomics. Molecular Biology and Evolution. 41:msae214.

Li X., Hu Y., Guo J., Lan J., Lin Q., Bao X., Yuan S., Wei M., Li Z., Man K., Yin Z., Han J., Zhang J., Zhu C., Zhao Z., Liu Y., Yang J., Nie J. 2022. A high-resolution climate simulation dataset for the past 540 million years. Sci Data. 9:371.

Matzke N.J. 2013. BioGeoBEARS: BioGeography with Bayesian (and likelihood) evolutionary analysis in R Scripts. R package, version 1.1.1. 1:2013.

Minh B.Q., Schmidt H.A., Chernomor O., Schrempf D., Woodhams M.D., von Haeseler A., Lanfear R. 2020. IQ-TREE 2: New models and efficient methods for phylogenetic inference in the genomic era. Molecular Biology and Evolution. 37:1530–1534.

Mitteroecker P., Collyer M.L., Adams D.C. 2025. Exploring Phylogenetic Signal in Multivariate Phenotypes by Maximizing Blomberg's K. Syst Biol. 74:215–229.

Mo Y.K., Lanfear R., Hahn M.W., Minh B.Q. 2023. Updated site concordance factors minimize effects of homoplasy and taxon sampling. Bioinformatics. 39:btac741.

Oliver P.M., Bauer A.M., Greenbaum E., Jackman T., Hobbie T. 2012. Molecular phylogenetics of the arboreal Australian gecko genus Oedura Gray 1842 (Gekkota: Diplodactylidae): Another plesiomorphic grade? Molecular Phylogenetics and Evolution. 63:255–264.

Oliver P.M., Sanders K.L. 2009. Molecular evidence for Gondwanan origins of multiple lineages within a diverse Australasian gecko radiation. Journal of Biogeography. 36:2044–2055.

Peel M.C., Finlayson B.L., McMahon T.A. 2007. Updated world map of the Köppen-Geiger climate classification. Hydrology and Earth System Sciences. 11:1633–1644.

Pepper M., Keogh J.S. 2021. Life in the "dead heart" of Australia: The geohistory of the Australian deserts and its impact on genetic diversity of arid zone lizards. Journal of Biogeography. 48:716–746.

Rannala B., Yang Z. 2017. Efficient Bayesian Species Tree Inference under the Multispecies Coalescent. Systematic Biology. 66:823–842.

Scotese C.R., Wright N. 2018. PALEOMAP Paleodigital Elevation Models (PaleoDEMS) for the Phanerozoic.

Singhal S., Grundler M., Colli G., Rabosky D.L. 2017. Squamate Conserved Loci (SqCL): A unified set of conserved loci for phylogenomics and population genetics of squamate reptiles. Molecular Ecology Resources. 17:e12–e24.

Skeels A., Sauquet H., Mast A., Weston P.H., Olde P.M., Jordan G.J., Carpenter R.J., Fenker J., Reynolds Z.K.M., Lemmon A.R., Lemmon E.M., Pichardo-Marcano F.J., Cardillo M. 2025. Paleobiome dynamics shaped a large Gondwanan plant radiation. Proceedings of the National Academy of Sciences. 122:e2502129122.

Skipwith P.L., Bauer A.M., Jackman T.R., Sadlier R.A. 2016. Old but not ancient: Coalescent species tree of New Caledonian geckos reveals recent post-inundation diversification. Journal of Biogeography. 43:1266–1276.

Skipwith P.L., Bi K., Oliver P.M. 2019. Relicts and radiations: Phylogenomics of an Australasian lizard clade with east Gondwanan origins (Gekkota: Diplodactyloidea). Molecular Phylogenetics and Evolution. 140:106589.

Smith S.A., Brown J.W., Walker J.F. 2018. So many genes, so little time: A practical approach to divergence-time estimation in the genomic era. PLoS ONE. 13:e0197433.

Steenwyk J.L., Iii T.J.B., Li Y., Shen X.-X., Rokas A. 2020. ClipKIT: A multiple sequence alignment trimming software for accurate phylogenomic inference. PLOS Biology. 18:e3001007.

Stein A., Gerstner K., Kreft H. 2014. Environmental heterogeneity as a universal driver of species richness across taxa, biomes and spatial scales. Ecology Letters. 17:866–880.

Tiatragul S., Brennan I.G., Broady E.S., Keogh J.S. 2023a. Australia's hidden radiation: Phylogenomics analysis reveals rapid Miocene radiation of blindsnakes. Molecular Phylogenetics and Evolution. 185:107812.

Tiatragul S., Skeels A., Keogh J.S. 2023b. Paleoenvironmental models for Australia and the impact of aridification on blindsnake diversification. Journal of Biogeography. 50:1899–1913.

Title P.O., Singhal S., Grundler M.C., Costa G.C., Pyron R.A., Colston T.J., Grundler M.R., Prates I., Stepanova N., Jones M.E.H., Cavalcanti L.B.Q., Colli G.R., Di-Poï N., Donnellan S.C., Moritz C., Mesquita D.O., Pianka E.R., Smith S.A., Vitt L.J., Rabosky D.L. 2024. The macroevolutionary singularity of snakes. Science. 383:918–923.

Yang Z. 2007. PAML 4: Phylogenetic Analysis by Maximum Likelihood. Mol Biol Evol. 24:1586–1591.

Yang Z. 2015. The BPP program for species tree estimation and species delimitation. Current Zoology. 61:854–865.

Zhang C., Mirarab S. 2022. Weighting by gene tree uncertainty improves accuracy of quartet-based species trees. Molecular Biology and Evolution. 39:msac215.

Zhang C., Zhao Y., Braun E.L., Mirarab S. 2021. TAPER: Pinpointing errors in multiple sequence alignments despite varying rates of evolution. Methods in Ecology and Evolution. 12:2145–2158.
